# Characterization of *NEB* mutations in patients reveals novel nemaline myopathy disease mechanisms and omecamtiv mecarbil force effects

**DOI:** 10.1101/2023.12.20.572678

**Authors:** Esmat Karimi, Mila van der Borgh, Johan Lindqvist, Jochen Gohlke, Zaynab Hourani, Justin Kolb, Stacy Cossette, Michael W. Lawlor, Coen Ottenheijm, Henk Granzier

## Abstract

Nebulin, a critical protein of the skeletal muscle thin filament, plays important roles in physiological processes such as regulating thin filament length (TFL), cross-bridge cycling, and myofibril alignment. Mutations in the nebulin gene (*NEB*) cause NEB-based nemaline myopathy (NEM2), a genetically heterogeneous disorder characterized by hypotonia and muscle weakness, currently lacking therapies targeting the underlying pathological mechanisms. In this study, we examined a cohort of ten NEM2 patients, each with unique mutations, aiming to understand their impact on mRNA, protein, and functional levels. Results show that truncation mutations affect *NEB* mRNA stability and lead to nonsense-mediated decay of the mutated transcript. Moreover, a high incidence of cryptic splice site activation was found in patients with splicing mutations which is expected to disrupt the actin-binding sites of nebulin. Determination of protein levels revealed patients with relatively normal nebulin levels and others with markedly reduced nebulin. We observed a positive relation between the reduction in nebulin and a reduction in TFL, and a positive relation between the reduction in nebulin level and the reduction in tension (both maximal and submaximal tension). Interestingly, our study revealed a duplication mutation in nebulin that resulted in a larger nebulin protein and longer TFL. Additionally, we investigated the effect of Omecamtiv mecarbil (OM), a small-molecule activator of cardiac myosin, on force production of type I muscle fibers of NEM2 patients. OM treatment substantially increased submaximal tension across all NEM2 patients ranging from 87-318%, with the largest effects in patients with the lowest level of nebulin. In summary, this study indicates that post-transcriptional or post-translational mechanisms regulate nebulin expression. Moreover, we propose that the pathomechanism of NEM2 involves not only shortened but also elongated thin filaments, along with the disruption of actin-binding sites resulting from splicing mutations. Significantly, our findings highlight the potential of OM treatment to improve skeletal muscle function in NEM2 patients, especially those with large reductions in nebulin levels.

## Introduction

Nemaline myopathy (NEM) is a rare, clinically and genetically heterogeneous disorder,^1^ characterized by hypotonia and muscle weakness.^2^ NEM is histopathologically defined by disorganization of the sarcomeric Z discs and the accumulation of nemaline bodies or rods in muscle fibers, which are aggregates of Z-disc and thin filament proteins.^3^ Moreover, NEM patients exhibit a shift towards type I fibers, a characteristic evident in both patient samples and mouse models of NEM.^4,5^ Studies indicate an estimated incidence of two cases per 100,000 live births, accounting for 17% of congenital myopathy cases.^6^ The muscle weakness ranges from severe to mild,^7-11^ and is most commonly congenital.^12,13^

Nemaline myopathies are associated with mutations in at least 13 genes: *ACTA1*, *NEB*, *LMOD3*, *TPM3*, *TPM2*, *TNNT1*, *TNNT3*, *CFL2*, *MYPN* (all encoding protein components of the muscle thin filament), *KBTBD13*, *KLHL40*, *KLHL41* (all likely involved in protein turnover in the muscle sarcomere via the ubiquitin-proteasome pathway), and *MYO18B.*^1,14^ These mutations can be either *de novo*, or inherited by autosomal dominant, or autosomal recessive patterns.^1^ Mutations in the *NEB* gene account for approximately 35% of nemaline myopathies,^15^ which are usually inherited recessively. However, a recent study identified the first dominantly inherited mutation in *NEB*, leading to a distal form of nemaline myopathy.^16^

Nebulin, a giant ∼800 KDa filamentous protein, is a crucial component of the thin filament in skeletal muscle.^17^ Single nebulin molecules span the thin filament, with their C-termini anchored in the Z-disk and N-terminal directed toward the pointed-end of the thin filament.^18^ In human muscle, nebulin primarily consists of 22 to 29 tandem super-repeats (SR).^19-21^ Each SR consists of seven simple repeats, each comprised of 31 to 38 amino acid residues, featuring a conserved sequence motif SDxxYK that is thought to be actin binding.^22^ Every nebulin simple repeat interacts with three neighboring actin subunits.^22^ Nebulin’s SRs interact with the troponin/tropomyosin regulatory complex via two contact sites between troponin T and nebulin, facilitated by troponin-binding motifs, including WLKGIGW and ExxK.^22^ Nebulin plays a critical role in various important processes in skeletal muscle such as maintaining Z-disk structure, myofibril alignment^23-25^ and crossbridge cycling.^26-31^ It has also been implicated in the regulation of thin filament length (TFL), as reduced levels of nebulin protein have been associated with shortened TFL in patients with NEB-related nemaline myopathy, as well as in nebulin-deficient mouse models, zebrafish, and chick skeletal myocytes.^30,32-39^

Despite investigating a number of therapeutic interventions including troponin activators such as Levosimendan,^40^ CK-20066260,^30^ dietary supplements^41^ and myostatin inhibitors,^42,43^ there are currently no effective therapies targeting the underlying pathological mechanisms for nebulin-based NEM (NEM2). It has been shown that Omecamtiv mecarbil (OM), a selective small-molecule activator of cardiac myosin (*MYH7)* that was initially developed as a treatment for heart failure,^44,45^, increases submaximal force production in type I skeletal muscles of Neb cKO mouse model^46^ and rat diaphragm.^47^ Considering the dominance of type I fibers in nemaline myopathy patients,^4,5^ it is expected that OM increases the force of type I fibers at submaximal activation levels in skeletal muscle fibers and so might improve the quality of life in nemaline myopathy patients.

Here we studied skeletal muscle tissue from a cohort of ten NEM2 patients with confirmed mutations in the *NEB* gene. We characterized these mutations by studying their effects on mRNA and protein levels as well as mechanical features of skeletal muscle. Our results revealed novel NEM disease mechanisms including disruption of actin binding sites by activation of cryptic splice sites. Furthermore, we showed that OM not only improves force production in slow skeletal muscle at submaximal activation levels, but that its effectiveness is greater in NEM2 patients relative to controls, with an inverse relation between nebulin level and OM-based tension increase.

## Results

### Mutation analysis using RNA sequencing

To assess the effects of *NEB* mutations on nebulin transcript level and mRNA processing, we performed RNAseq analysis on skeletal muscle biopsies of 10 NEM2 patients (supplementary Table 1 presents clinical details) and as controls, three healthy individuals who did not exhibit any muscle-related conditions. Patient 144 had a homozygous deletion of exon 55, whereas the remaining patients exhibited compound heterozygous mutations, including splice site mutations, truncations, frameshifts, duplications, insertions, or deletions (mutation details in Table 1). A summary of all RNA-seq findings (explained below in detail) can be found in Supplementary Table 2.

**Table 1.** Genetic information of patients with NEM2.

| Patient ID | Mutation Details in NEB gene |  | Mutation consequence | Mutation type | Clinical significance | Mutation site on nebulin protein |
| --- | --- | --- | --- | --- | --- | --- |
| 2385 | 1 | exon 32 c.3252_3255+3delTGACGTA | Results 7 bp deletion | Deletion | Likely pathogenic | SR3 R6,R7 |
|  | 2 | 2.7 kb deletion exon 77 g.152,469,299 in intron 77 g.152,471,995 in intron 76 | Results 2697 deletion including exon 77 | Deletion | - | SR14 R7 / SR15 R1,R2,R3 |
| 2296 | 1 | exon 112 c.17654G>A | Results generation of stop codon in exon 112 | Truncation | Pathogenic | SR24 R2/R3 |
|  | 2 | exon 175 c.24771delT | Results deletion in exon 175 leading to stop 19 bp into exon 176 | Frameshift | Pathogenic/Likely pathogenic | M234/M235 |
| 2486 | 1 | exon 61 c.8425 C>T | Results generation of stop codon in exon 61 | Truncation | Pathogenic/Likely pathogenic in LOVD | SR10 R7 / SR11 R1,R2,R3 |
|  | 2 | exon 171 c.24317 T>A | Results generation of stop codon in exon 171 | Truncation | Pathogenic/Likely pathogenic in LOVD | M230/M231 |
| 3424 | 1 | exon 85 c.13059+5G>A | Results point mutation in intron 85 | Intronic point mutation | Likely pathogenic | SR16 R7 / SR17 R1,R2,R3 |
|  | 2 | exon 169 c.24218C>A | Results generation of stop codon in last codon of exon 169 | Truncation | VUS/pathogenic in LOVD | M228/M229 (Z-disk) |
| 4001 | 1 | exon 32 c.3255+1G>A | Results donor splice mutation in junction of exon-intron 32 | Splicing | Pathogenic | SR3 R6/R7 |
|  | 2 | exon 110 c.17501_17502delinsC | Results insertion in exon 110 leading to stop codon 19 bp into exon 111 | Frameshift | Pathogenic in LOVD | SR23 R7/SR24 R1 |
| 2622 | 1 | exon 80 c.12018+1G>A | Results donor splice site mutation in junction of exon-intron 80 | Splicing | Pathogenic/Likely pathogenic | SR15 R6/R7 |
|  | 2 | exon 172 c.24458_24461dupAGAT | Duplication of AGAT in exon 172 leads to stop codon at the end of exon | Frameshift | Pathogenic/Likely pathogenic | M231/M232 |
| 4526 | 1 | exon 109 c.17262G>A | Generation of stop codon in the middle of exon 109 | Truncation | Pathogenic | SR22 R6/R7 |
|  | 2 | exon 171 c.24318_24319insAA | Results insertion at the start of exon 171 leading to stop codon in exon 172 | Frameshift | Pathogenic | M229/M230 (Z-disk) |
| 144.0 | 1<br>2 | Homozygous exon 55 deletion | Results in-frame 2502 deletion including exon 55 | Deletion | Pathogenic | SR9 R5,R6 |
| 180.0 | 1 | Four copy gain within the NEB triplicate repeat region | 10 TRI repeats instead of 6 | Duplication | Pathogenic | [SR16 R3,R4,R5] – [SR21 R7 / SR22 R1,R2,R3] |
|  | 2 | exon 157 c.22936C>T (p.Arg7646*) | Results generation of stop codon in exon 157 | Truncation | Pathogenic | M216/M217 |
| 151.0 | 1 | exon 10 c.822+1G>A | Results donor splice site mutation at the junction of exon-intron 10 | Splicing | Likely pathogenic | M5 |
|  | 2 | and exon 30 c.3042+3_3042+6del | Results 4 bp deletion of intron 30 | Intronic deletion | VUS | R5 of SR2 |

Several regions in *NEB* are known to undergo alternative splicing (the central SR region exons 63-66, 82-105, and 143/144; the Z-repeat region exons 166-177) and it has been proposed that mutations in these regions may have a relatively mild pathogenic effect compared to mutations elsewhere in the gene.^20,21,48^ We performed alternative splicing analysis and studied RNA processing in each sample. Findings confirm alternative splicing in the Z-repeat exons 166-177, as well as the SR exons 143/144 and 82-105, as shown by their PSI values below 1.0 (Fig. 1A). Moreover, we observed the exclusion of specific exons in several patients, which could be explained by the known mutations present in those individuals. For instance, patient 144, who had a homozygous deletion of exon 55, exhibited a PSI of 4% for that exon, confirming the near absence of this exon in the majority of nebulin mRNA molecules in this patient (Fig. 1A). Similarly, patient 2385, with a deletion of exon 77 on one of their *NEB* alleles, displayed a PSI of 46% for exon 77, which is close to the expected 50% PSI. Additionally, in patient 151, exon 30 was skipped in 85% of the nebulin transcripts. This patient carried a 4-base pair deletion in intron 30 (c.3042+3_3042+6del), classified as a variant of uncertain significance (VUS) in the ClinVar database. The splice site analysis further supported the pathogenic nature of this intronic mutation.

**Figure 1.**
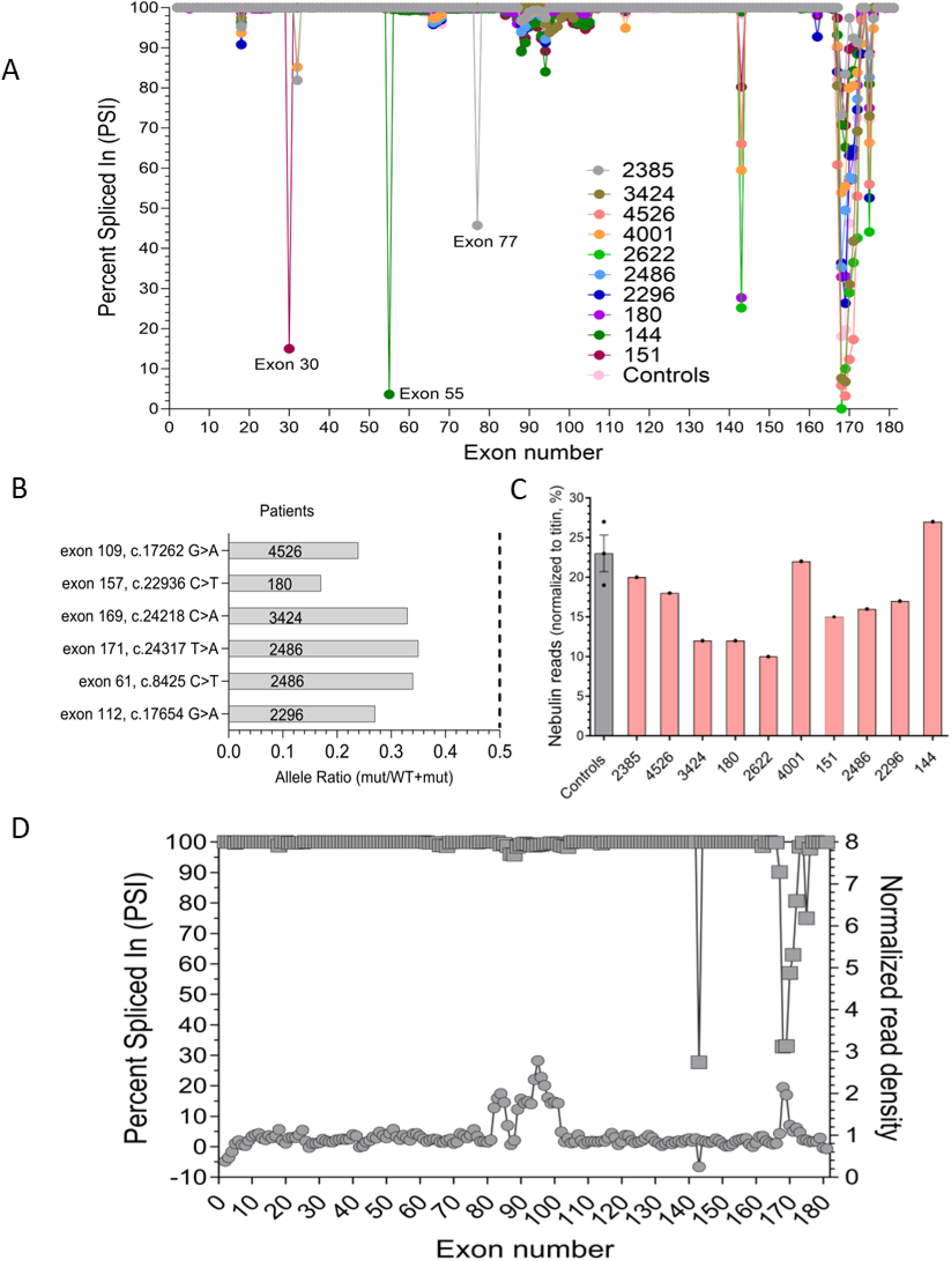
Effect of mutations on *NEB* mRNA. A) Alternative splicing pattern of *NEB* gene in NEM2 patients and controls. Percent spliced-in (PSI) values range from 0% (excluded in all transcripts) to 100% (included in all transcripts). B) Allelic imbalance analysis for truncation mutations of NEM2 patients. The bars represent the expression ratio of the mutated allele to WT allele. The dotted line indicates the expected ratio if both alleles are expressed equally. C) Read counts of NEB gene normalized by *TTN* reads. D) Expression analysis of patient 180. The left Y-axis shows the PSI of *NEB* exons (square). To differentiate the copy gains in the triplicated region of *NEB*, the normalized read density of each exon is shown on the right Y-axis (circle), revealing a higher read density for exons in the triplicated region.

Truncation mutations are frequently observed in NEM2 patients (Table 1). To better understand the impact of truncation mutations on the stability of *NEB* mRNA in these patients, we conducted an allelic imbalance analysis. This involved summing the reads based on their position and comparing the number of mutated alleles to wild-type alleles at the site of the truncation mutation. A lower ratio of mutant to WT alleles would suggest that the mutated transcript undergoes nonsense-mediated decay (NMD).^49^ Our findings, shown in Fig. 1B, reveal allelic imbalance in patients 2296, 2486, 3424, 4526, and 180, all of whom harbor truncation mutations. This supports the notion that truncation mutations in these patients affect *NEB* mRNA stability and lead to NMD of the mutated transcript.

To evaluate the overall transcript level of nebulin, we normalized nebulin read counts to titin, to account for varying ratios of muscle fibers contained within each biopsy sample and for differences in library size. Results depicted in Fig. 1C show that nebulin transcript levels of patients are either relatively normal (patients 2385, 4001, 4526 and 144) or are reduced (patients 3424, 180, 2486 and 2622). Varying degrees of nebulin transcripts levels might be due to different NMD efficiency or skipping of exons carrying stop codon (Supplementary Table 2).

Additional analysis was also performed on patient 180. According to genetic testing, one of the *NEB* mutations in this patient is a four-copy gain in the triplicate region of nebulin. The triplicated (TRI) region of nebulin normally consists of 3 groups of exons (exons 82-89, 90-97, 98-105) each representing 2 SRs for a total of 6 SRs.^50^ Four copy-gain results in ten TRI repeats, which if transcribed and translated fully would result in 8 more SRs in nebulin in this patient.^50^ To investigate the TRI expression level, we normalized read counts of each exon in this patient by exon length and then measured the ratio of normalized read density of all exons relative to controls. Fig. 1D illustrates the results. The normalized read density of exons within the triplicate region (82-105) varied somewhat but was around twice as high as controls in most exons. Some exons were closer to control read density (exons 87,88 and 102-105) while others were increased about three-fold (exons 94-97). Mapping of RNA-seq reads carries uncertainty due to the repetitive nature of the region. Overall, increased normalized read density versus controls would implicate an addition of 2859 bases versus controls, which would equate to an addition of 28 simple repeats (two TRI copies or four SRs) (Supplementary Table 3). Thus, in contrast to the genetic testing report of this patient which shows four more TRI repeats at DNA level, analysis at the transcription level shows only two additional TRI repeats. Furthermore, analysis of allelic imbalance of this patient’s truncation mutation in exon 157 indicated that transcripts originating from the other allele accounted for only 17% of all transcripts (Fig. 1B). This suggests degradation of transcript of this allele by NMD.

Partial intronic inclusion and activation of cryptic splice site was consistently observed in all patients with mutations at donor splice sites (variants 2385, 4001, 2622, 151, and 3424), which resulted in atypical alternative splicing patterns, as detailed in Fig. 2A. Cryptic splice sites are normally inactive or utilized at low levels unless prompted by mutations near authentic splice sites.^51^ While it is commonly believed that the activation of cryptic splice sites may play a role in a wide range of genetic diseases,^52^ the concept of reversing aberrant splicing through the targeted activation of cryptic splice sites has been suggested as a potential therapeutic strategy.^53^ The activation of cryptic splice site in NEM2 patients led to either in-frame (4001, 2622 and 3424) or out-frame (2385 and 151) intronic inclusion. In-frame intron inclusion could lead to production of nebulin transcripts with increased repeat lengths. Moreover, the transcripts produced by cryptic splice sites showed different percentages in NEM2 patients (Figure 2A and Supplementary Table 2). The percentage of transcript produced by cryptic splice site activation, in-frame or out-frame intronic inclusion and percentage of other transcript isoforms determine the different effects of cryptic splice site activation on the protein level. For instance, we detected partial inclusion of intron 32 in patients 2385 and 4001, which would result in an addition of 13 amino acids to repeat 7 of SR3. In patient 4001, this transcript version is in frame and constitutes around half of all nebulin isoforms, and it could have a significant impact on nebulin structure due to increasing repeat length. Interestingly, our results showed that intron inclusion in this patient would break up an actin binding motif and add unstructured sequence into sections of nebulin which are predicted to be α-helical by ColabFold^54^ (Fig. 2B and Supplementary Fig. 1A). This would lead to a mismatch between thin filament and nebulin binding sites which need to be identically spaced. This finding provides insight into the underlying pathomechanism of splicing site mutations in NEM2 patients.

**Figure 2.**
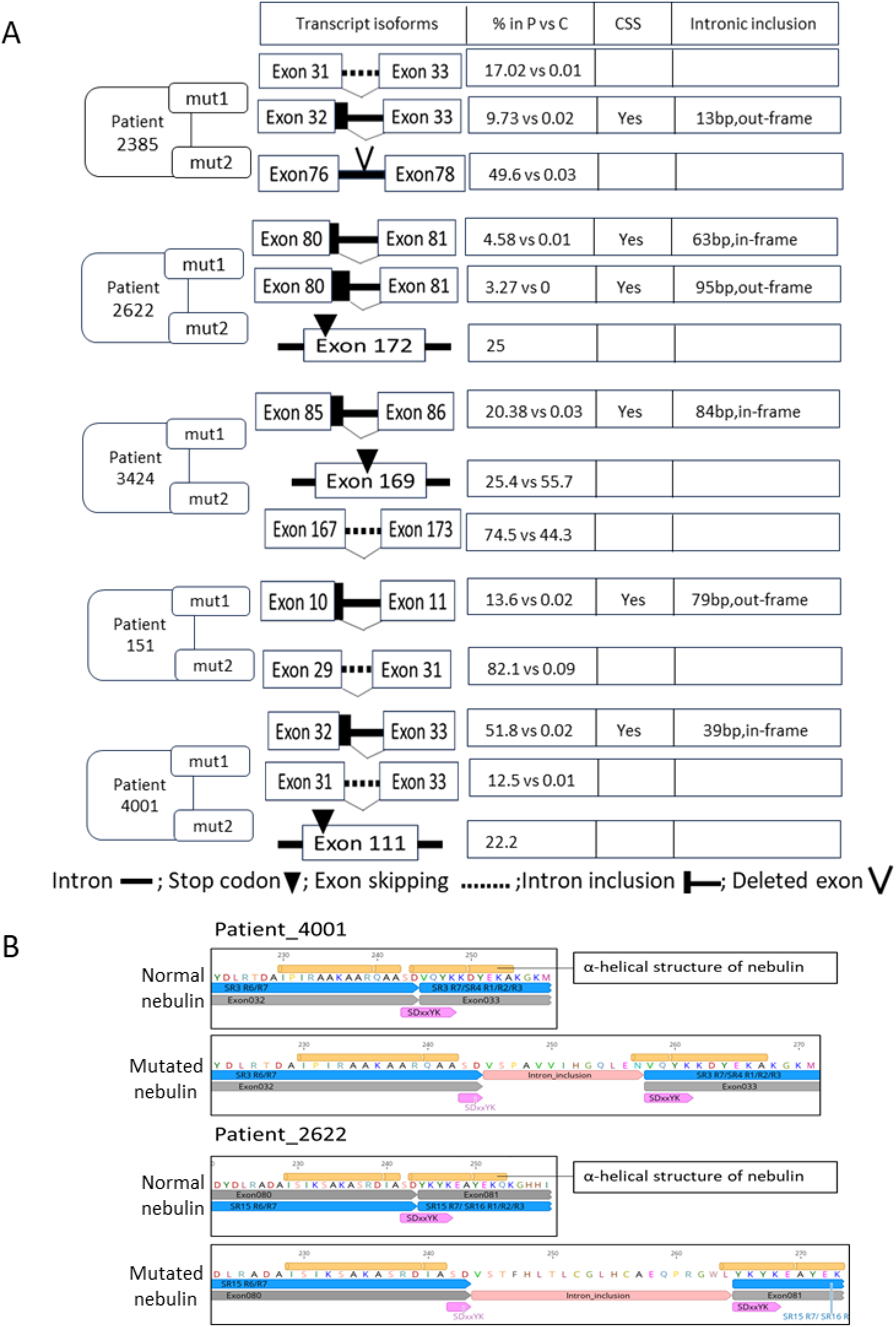
Effect of *NEB* splice site mutations on transcript and protein. A) In all patients with splice site mutations, cryptic splice sites are activated. For each patient and each mutation, transcript isoforms, percentage of transcript isoform in patient versus controls (% in P vs C), cryptic splice site activation (CSS) and intronic inclusion are indicated. The mutation (mut1 and mut2) details of each patient can be found in Table 1. The shown transcript isoforms are only those impacted by mutation. B) Prediction of nebulin structure at the site of splicing mutations. In patient 4001 (top), splicing mutation results in 39 bp in-frame inclusion of intron 32 and consequently 13 additional amino acids in super-repeat 3; in patient 2622 (bottom), 63 bp in-frame inclusion of intron 80 results in 21 additional amino acids in super-repeat 15. In both patients, intronic inclusion leads to disrupted actin-binding sites on nebulin. The predicted nebulin α-helix structure is in yellow, super-repeat region is in blue, actin-binding motif is in pink and exon is in grey (bottom).

Partial intron 32 inclusion in patient 2385 accounts for only 10% of transcripts because intron inclusion is out-of-frame due to the 7bp deletion (Fig. 2A and Supplementary Table 2) and leads to a stop codon and subsequent degradation by NMD. This might also explain why the ratio of the isoform with skipping of exon 32 relative to the isoform with intron inclusion is higher in patient 2385 (splice site deletion) than patient 4001 (splice site substitution). Other significant intron inclusion events occur in patient 2622, where inclusion of a portion of intron 80 adds 21 amino acids to repeat 7 of SR15. Like patient 4001, this intron inclusion event would add an unstructured sequence in the middle of nebulin’s actin binding motif (Fig. 2B and Supplementary Fig. 1B). In patients 3424 and 151, cryptic donor splice sites downstream of the mutation were activated, but transcripts created from these sites would most likely be subject to NMD due to the presence of stop codons (Supplementary Fig. 2).

In summary, disease mechanisms that were revealed include degradation of nebulin transcript by NMD and exon skipping, and activation of cryptic splice sites that disrupt nebulin binding sites to the thin filament. Additionally, the obtained results reveal for the first time that in some patients, longer than normal transcript can be produced.

### Nebulin protein expression

To assess the impact of *NEB* mutations on nebulin protein levels, we used sodium dodecyl sulfate-agarose gel electrophoresis (Fig. 3A and B) and western blot (Fig. 3C) techniques on lysates from muscle biopsies from both NEM2-patients and controls. Nebulin protein levels were normalized using myosin heavy chain (MHC) as a reference for protein loading. Previous studies have consistently reported a nebulin:MHC ratio of ∼0.05-0.07across various skeletal muscle types and vertebrate species.^55-57^ In line with this, our study revealed a nebulin:MHC ratio in control samples of 0.068 (Fig. 3B). When analyzing patients’ samples, results segregated into two groups. Patients 3424, 4526, 180, and 2385 displayed nebulin:MHC ratios of 0.07, 0.07, 0.05, and 0.06, respectively. These ratios are all within the range of nebulin:MHC ratios observed in the control group (Fig. 3B-left). However, patients 4001, 2622, 2486, 144, and 151 displayed lower ratios, hovering around 0.03, except for patient 2296, where nebulin was undetectable on protein gels (Fig. 3B-left). When results were grouped accordingly, a significant reduction in nebulin expression was found in the patients with low expression (Fig. 3B-right). Furthermore, to determine if the patients expressed full-length nebulin, we utilized Western blotting with antibodies targeting the N-terminal and C-terminal ends of nebulin. The results revealed the presence of full-length nebulin in all patients (Fig. 3C). It should be noted that this full-length nebulin could potentially be mutated (such as those with in-frame deletions) or non-mutated.

**Figure 3.**
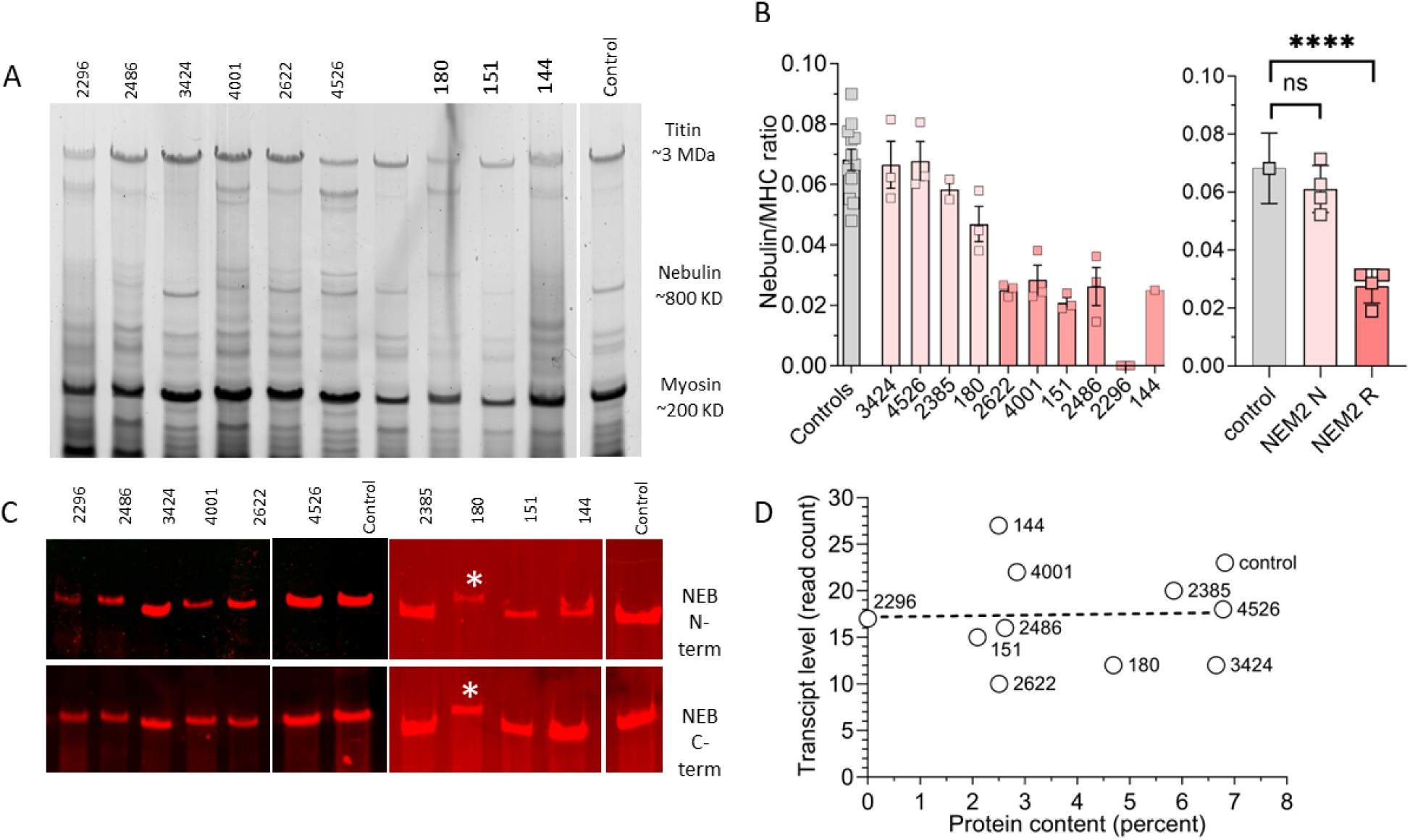
Protein expression analysis. A) Agarose gel electrophoresis reveals detectable nebulin in all patients, except for patient 2296. B) (left) Nebulin quantification is expressed as nebulin/MHC ratio. Patients 3424, 2385, 4526, and 180 have nebulin levels similar to controls, while patients 2622, 4001, 151, 2486, 2296, and 144 have reduced nebulin levels. B) (right) Mean values per biopsy grouped as controls, NEM2 patient with normal nebulin (NEM2 N) and reduced nebulin (NEM2 R) expression and differences between groups analyzed with a nested one-way ANOVA. Asterisks indicate a significant difference between NEM2 R and controls. \**P*<0.05, ** *P* <0.01, *** *P* <0.001, **** *P* <0.0001 and ns indicates no significant difference. C) Western blot analysis using antibodies against the N-terminal and C-terminal ends of nebulin, targeting the M1-M3 and SH3 domains, respectively. Asterisk reveals slower migrating nebulin in patient 180. D) Transcript level vs protein level in controls and patients shows lack of a correlation (slope of shown regression line not different from zero, *P*=0.9). Note that in B left) samples were analyzed independently 3 times (technical replicates) and that results from 3 controls were pooled.

The mobility of nebulin on agarose gel was similar for most patients, except for patient 180, who displayed visibly reduced nebulin mobility (see asterisk in Fig. 3C), suggesting a larger size of nebulin. This observation is consistent with this patient’s mutations and aligns with the RNA-seq results, which showed higher read density of exons in the triplicated region. As discussed above, according to genetic testing, one of the *NEB* mutations in patient 180 involves a four-copy gain in the triplicate region of nebulin. However, RNA-seq analysis showed transcripts produced by this allele contain 2858 additional base pairs (Supplementary Table 3) which accounts for two TRI copies and, thus, 4 more SRs. At the protein level this is expected to result in a nebulin that is 108 kDa (4x 27 KD) larger than normal, consistent with the reduced mobility of the nebulin band on the agarose gel for this patient. Additionally, as described earlier, the other allele of patient 180 carries a truncation mutation in exon 157, and accounts for only 17% of all transcripts, indicating its likely degradation through the NMD mechanism (Fig. 1B). It is noteworthy that protein gels do not reveal a band that is associated with this allele (Fig. 3A and 3C) implying another mechanism to prevent translation of transcript that carries the truncation mutation.

The transcript and protein data for each patient make it possible to evaluate if transcript and protein levels are correlated. Fig. 3D reveals that the nebulin transcript’ read counts for patients with low nebulin protein levels are not different from those with normal protein and the linear regression line has a slope that is not different from 0 (*P*=0.9), i.e., there is no correlation between nebulin transcript and protein levels.

### Effect of *NEB* mutations on TFL

Considering that nebulin plays a role in TFL specification,^57^ we investigated the effect of *NEB* mutation on TFL. To analyze muscle biopsies, we cut longitudinal sections and stained them with fluorescently labeled phalloidin to mark the actin filaments. TFL was measured according to the detailed procedures outlined in the Methods section. The results of all patients, except patient 180 (for reasons, see below) are presented in Fig. 4A. The TFL measurement revealed that patients with normal levels of nebulin (3424, 4526, and 2385) displayed mean thin filament lengths like those of the control group. In contrast, patients with reduced levels of nebulin (2622, 2296, 151 and 2486) exhibited a significant decrease in TFL compared to the controls (Fig. 4B). Linear regression analysis between TFL and nebulin content reduction revealed a significant negative relation (*P*=0.003) (Fig. 4C). These findings support an important role of nebulin in determining TFL, as demonstrated by shorter TFL when nebulin protein levels are reduced.

**Figure 4.**
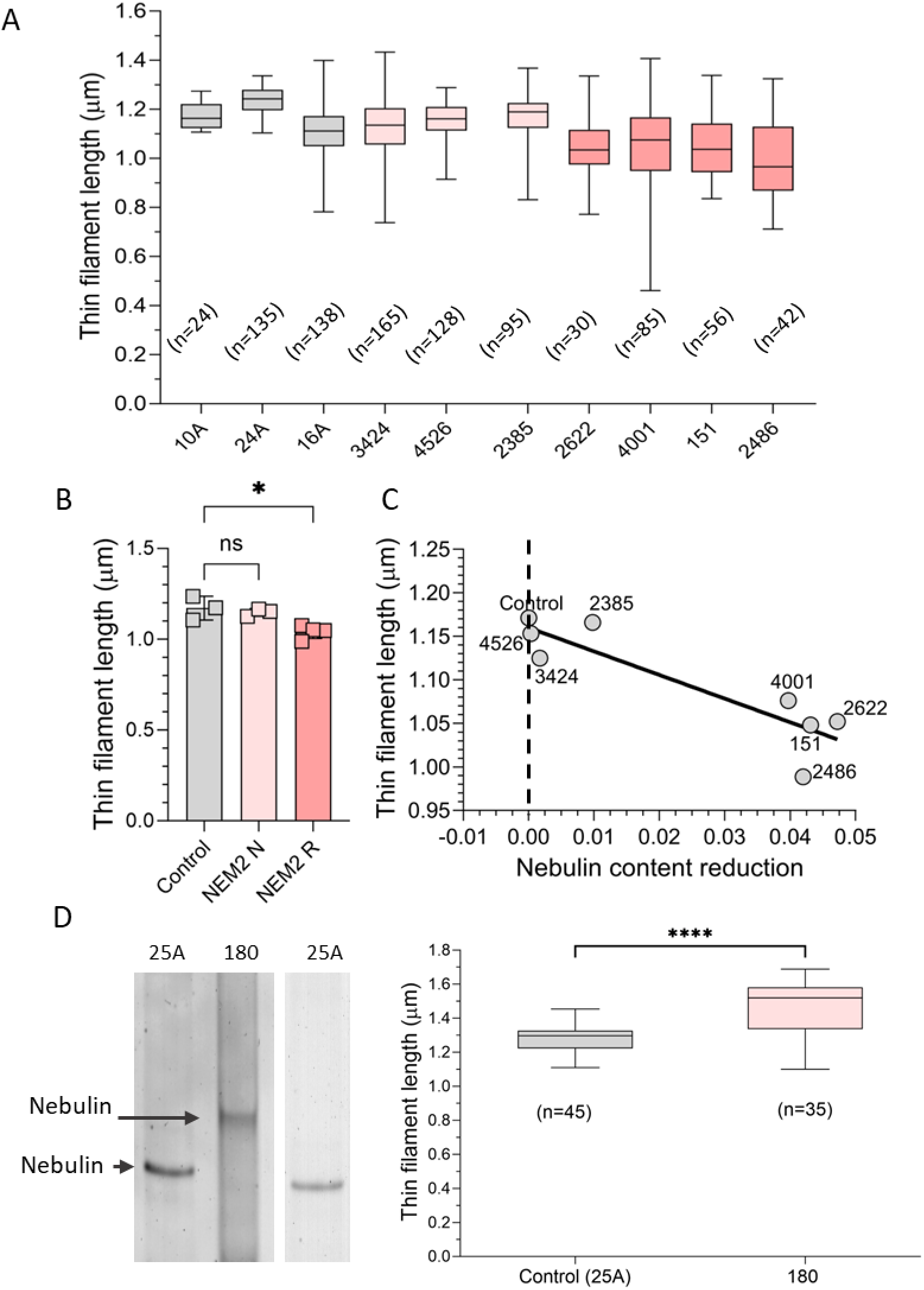
Thin filament length (TFL) in muscle fibers from NEM2 patients and controls. A) TFL was measured based on phalloidin staining of actin filaments of NEM2 patients (except patient 180) and controls. Measurements were obtained in the 2.8-3.2 mm sarcomere length range. B) Mean TFL per biopsy, grouped with NEM2 biopsies segregated according to their nebulin expression level (N for normal and R for reduced, see Figure 3 for details). One-way ANOVA analysis reveals significantly reduced TFL in NEM2-R samples. C) Mean TFL vs. nebulin content reduction (nebulin:MHC ratio in controls minus in patients). The slope of the linear regression line is significantly different from zero (*P*=0.003). D) (Left) electrophoretic analysis of control and patient 180 samples on a 1% agarose gel reveals reduced mobility of nebulin in patient 180. (Right) TFL measurements (sarcomere length range 3.5-4.0 mm) show longer TFL in patient 180 (1.469 mm) than control (1.275 mm). Data are presented as box and whisker plots. Asterisks indicate a significant difference between NEM2 R and controls. \**P*<0.05, ** *P* <0.01, *** *P* <0.001, **** *P* <0.0001 and ns indicates no significant difference.

RNA-seq experiments suggested a larger nebulin in patient 180. This was confirmed by agarose gel electrophoresis that showed a nebulin band with less mobility indicating that a larger nebulin protein is expressed (Fig. 4D-left). Therefore, a separate study was performed where bundles from patient 180 and one control biopsy was stretched to long sarcomere lengths (3.5-4.0 µm). TFL measurement showed that patient 180 had significantly longer thin filaments than controls (Fig. 4D-right).

### Effect of *NEB* mutations on tension

It is well established that nebulin is important for normal contraction,^33^ therefore, we also investigated Ca^2+^-activated force in single fibers. Frozen biopsies were thawed and membrane-permeabilized, and single fibers were carefully dissected and mounted in an apparatus for subsequent force measurements. These initial measurements were conducted at two activation levels: maximal (pCa(-log[Ca2+]) 4.0) and submaximal calcium activation (pCa 6.75). After the mechanical study, fibers were fiber-typed (see Methods for details).

Several of the patients’ biopsies expressed mainly slow fibers (type I). Since we decided to study the effect of the slow myosin activator OM^58^ (see below), the results of tension production are focused on type I fibers. The control group’s type I fibers exhibited robust levels of maximal tension, averaging around 160 mN/mm². In contrast, the results from NEM2 patients displayed considerable variation. Several patients with relatively normal levels of nebulin (3424, 4526, 2385) showed the highest tensions, reaching approximately two-thirds of the control values. Conversely, patients with reduced nebulin levels experienced the lowest tensions, reaching only around one-third of the control values (Fig. 5A-left). Plotting the obtained tension values against nebulin protein reduction (Fig. 5A-right) revealed a significant negative relationship (*P*=0.004). Similarly, submaximal activation resulted in reduced tensions among patients (Fig. 5B-left), and a negative relation (*P*=0.0004) was identified between submaximal tension and nebulin protein reduction (Fig. 5B-right). Thus, the extent to which *NEB* mutations depress tension appears to be strongly dependent on the reduction in nebulin protein content.

**Figure 5.**
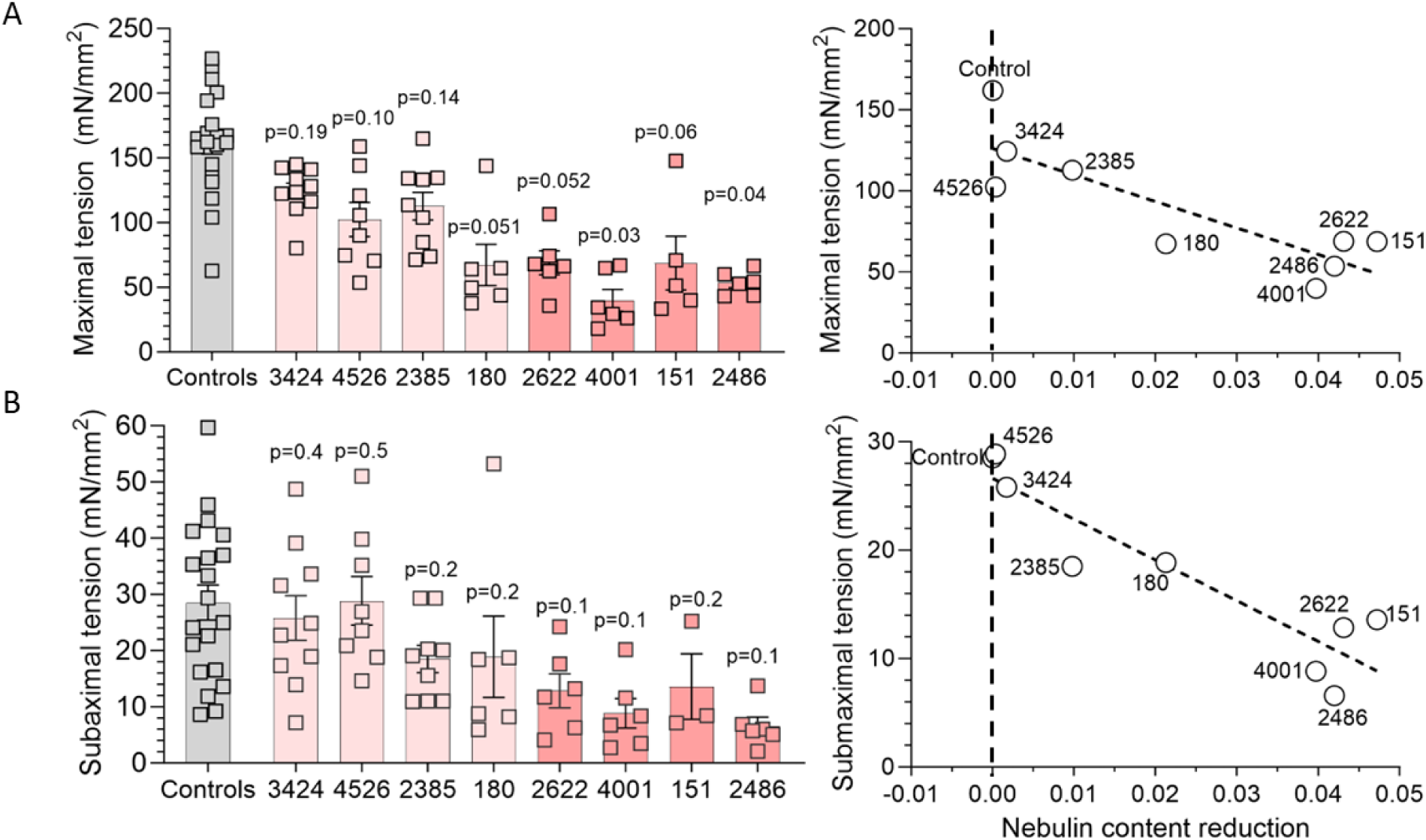
Tension production in control and NEM2 patients. Maximal tension at pCa 4.0 (A) and submaximal tension at pCa 6.75 (B) of type I fibers in control and NEM2 fibers. Left: results from individual fibers; right: mean tension vs nebulin protein content. Both maximal and submaximal tensions negatively correlate with nebulin content reduction (NEB/MHC ratio in controls minus in patients). Note that each data point in A left and B left reflects one fiber, and in B and D right the mean of all fibers per biopsy. Data were analyzed by linear regression analysis and obtained *P*-value for slopes are 0.004 (A) and 0.0004 (B). *P* values on each bar in A and C left are obtained by nested t-test analysis to compare tension between each patient and controls.

### Enhancing force development with OM

To assess the potential of OM in enhancing muscle force among patients with NEM2, patient fibers were activated in solutions with varying levels of calcium (pCa 8 to pCa 4) while being exposed to either 0.5 μM OM or vehicle. By analyzing the relationship between force and calcium concentration, we determined several parameters, including maximal tension (at pCa 4.0), submaximal tension (at pCa 6.75), pCa50 value (which represents the calcium concentration at half-maximal tension), and the Hill coefficient (nH), a measure of the cooperativity of activation.

Experiments revealed that OM did not affect type II fibers (Supplemental Fig. 3). This finding aligns with previous research indicating that OM selectively affects fiber types expressing the *MYH7* gene, such as cardiac muscle and type I skeletal muscle fibers.^46,47,59^ No significant effect on maximal tension was observed when comparing OM-treated type I fibers with those treated with vehicle, in either patients or controls (Supplemental Fig. 4). This suggests that 0.5 μM OM treatment does not reduce the force deficit observed at maximal activation levels in these patients, which is consistent with earlier studies.^46,47^

In type I fibers, the force-pCa plots exhibited a notable leftward shift in fibers treated with 0.5 µM OM compared to the control group (vehicle) (Fig. 6A). This shift was characterized by an increased pCa50 value (Fig. 6B) and a decrease in nH (not shown). Moreover, comparing the shift in pCa50 between control subjects and NEM2 patients, we observed a more pronounced effect in the NEM2 group (Fig. 6C). These findings indicate that OM enhances calcium sensitivity, with a greater impact observed in NEM2 biopsies. Additionally, submaximal tension measurements demonstrated significant OM-based increases (87-318%) in both control and NEM2 biopsies (Fig. 6D and 6E). Notably, the tension increase was more prominent in NEM2 patients and showed an inverse relationship with nebulin protein content (Fig. 6F) i.e., the increase in tension is highest in patients with the lowest level of nebulin.

**Figure 6.**
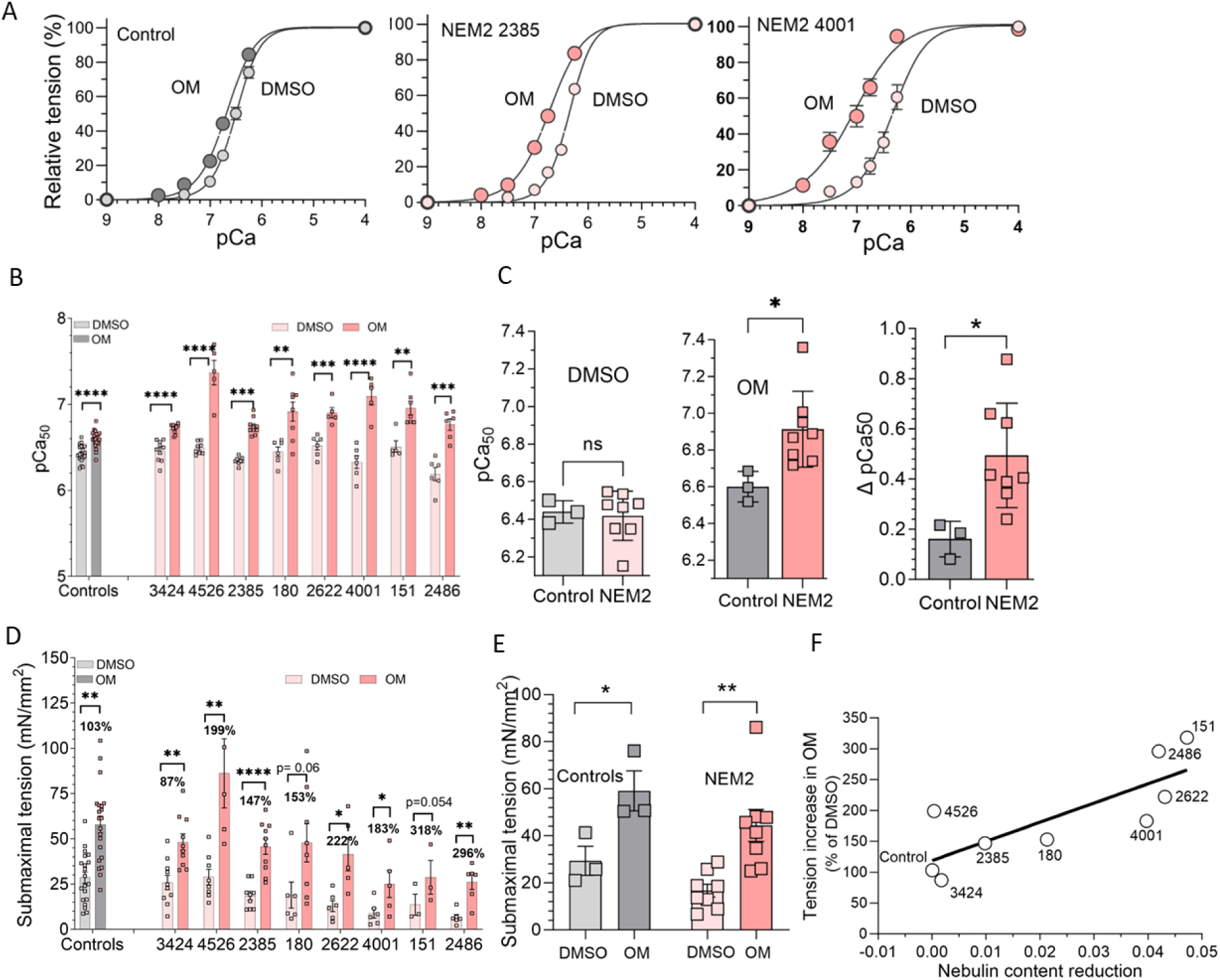
Effect of omecamtiv mecarbil (OM) on calcium-sensitivity and submaximal force. A) Example force-pCa curves in control and NEM2 patient with normal nebulin level (2385) or reduced nebulin (4001). OM left shifts the curves. B) Summarized pCa50 data of all biopsies revealing that OM increases pCa50 in all cases (each data point represents a separate single fiber). C) pCa50 of the control and NEM2 biopsies grouped and analyzed in an unpaired t-test showing no difference between groups in DMSO (left) but a significantly higher pCa50 in OM in NEM2 patients (middle) and consequently a larger DpCa50 in NEM2 patients (right). D) Submaximal tension (pCa 6.75) of control and NEM2 fibers is increased by OM in all patients and controls. E) t-test analysis shows that summarized tension of control and NEM2 fibers are significantly increased by OM. F) Tension increase in OM scales with nebulin content reduction (NEB/MHC ratio in controls minus in patients) (p-value of regression line: 0.01). Asterisks indicate a significant difference between DMSO and OM-treated fibers and ns indicates no significant difference. \**P*<0.05, ** *P* <0.01, *** *P* <0.001, **** *P* <0.0001

### Effect of OM on rate of tension development (*ktr*) and dynamic stiffness

Utilizing membrane-permeabilized single muscle fibers we studied the effect of OM on *Ktr at* maximal (pCa 4) and submaximal (pCa 6.75) activation levels. A series of step length changes was also imposed and the ensuing force transients were fitted to a non-linear distortion recruitment model, to extract the number of attached cross-bridges (ED) and their attachment and detachment rates.^60^ To keep the workload manageable, we used fibers from controls and two patients: patient 2385 with normal nebulin and patient 2486 with reduced nebulin. Experimental details and typical force recording are provided in the Methods section and Supplementary Fig. 5.

Findings show OM leads to a reduction in *ktr* for controls and both patients at both pCa 4 and pCa 6.75 (Fig. 7A). This decrease can be attributed to the extended duration of the strongly bound state of myosin heads in the presence of OM, culminating in slower cross-bridges cycling.^61^ Moreover, at pCa 4, OM treatment does not alter ED^62-65^ (Fig. 7B-left) but at pCa 6.75 OM increases ED in controls and patients (Fig. 7B-right).

**Figure 7.**
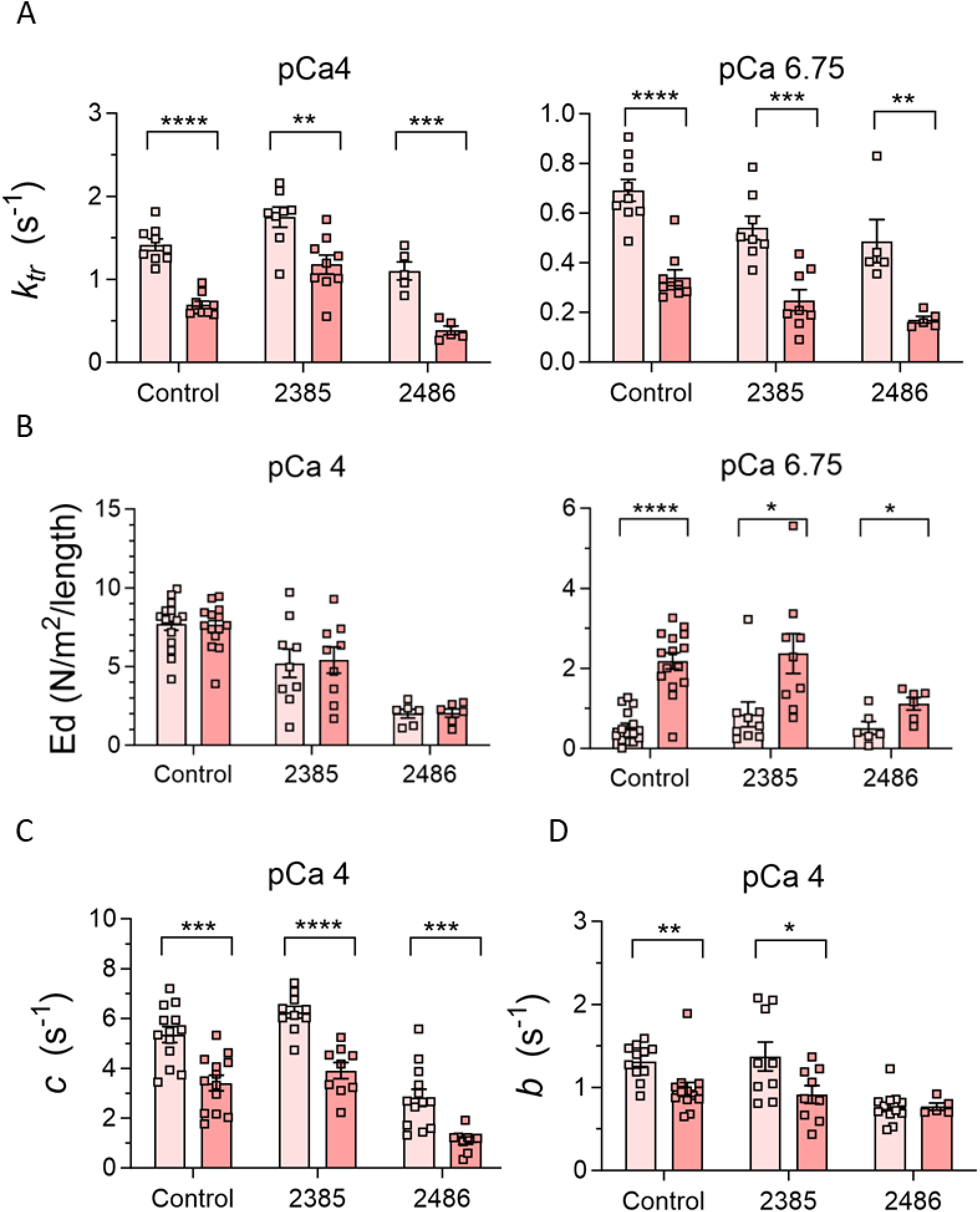
Effect of Omecamtiv mecarbil (OM) on rate of force redevelopment (*ktr*) and dynamic stiffness and crossbridge kinetics. A) OM results in lower *ktr* in controls and patients 2385 (normal nebulin) and 2486 (reduced nebulin) at both pCa4 (left) and pCa6.75 (right), B) OM treatment does not affect dynamic stiffness (Ed) at pCa4 (left) but increases it significantly at pCa6.75 in controls and both patients (right), C) OM treatment lowers the detachment rate of cross-bridges (*c*) at pCa4 in controls and both patients, D) OM treatment lowers the attachment rate of cross-bridges (*b*) at pCa4 in controls and patient 2385. Asterisks indicate a significant difference between DMSO and OM-treated fibers and ns indicates no significant difference. * *P* <0.05, ** *P* <0.01, *** *P* <0.001, **** *P* <0.0001

Finally, we measured detachment (*c*) and attachment (*b*) rate of cross-bridges at pCa 4. (Due to the low force levels at pCa 6.75, only a limited data set could be obtained, see Supplementary Fig. 6). As shown in Fig. 7C, OM-treated fibers exhibited significantly decreased *c* in controls and both patients. However, OM treatment resulted in reduced *b* in only controls and patient 2385 and in patient 2486 did not show an effect (Fig. 7D).

## Discussion

We investigated a cohort of NEM2 patients, each with distinct mutations, and studied their impact on mRNA, protein, and functional levels. We found that truncation mutations reduce *NEB* mRNA stability and lead to NMD of the mutated transcript. Additionally, a high occurrence of cryptic splice site activation and intronic inclusions was detected. In several NEM2 patients, intron inclusions added an unstructured region to nebulin’s simple-repeats, which is expected to disrupt the actin-binding sites of nebulin. At the protein level, the expression level of nebulin varied in NEM2 patients, between close to normal and very low, and no correlation between nebulin transcript and protein levels were found. Force and TFL were reduced in NEM2 patients (with the exception of one patient exhibiting longer thin filaments), and the effect was most severe in patients with the lowest nebulin levels. Finally, OM increased submaximal force levels and the effect was highest in patients with the lowest nebulin level. Below, we elaborate on these results.

### Mutation analysis and effect on nebulin expression

Based on quantification of nebulin protein levels, patients were divided into two groups: one with approximately normal levels of nebulin protein (average nebulin/MHC ratio of 0.06) and one with reduced levels (average nebulin/MHC ratio of 0.03). The overall *NEB* transcript level was not correlated with nebulin protein levels (Fig. 1C). This discrepancy is consistent with other genome-wide studies that also have shown that the correlation between levels of mRNA and protein is generally poor, hovering around 40% explanatory power across many studies^66,67^. This may be due to several factors, such as variations in translational rates or protein turnover, as well as the presence of post-transcriptional/translational mechanisms that regulate nebulin expression.

Previous studies have proposed that partial SR deletion is not tolerated in nebulin^30,57,68^ because it causes a structural mismatch between nebulin actin binding sites and the thin filament, and this prevents nebulin incorporation in the thin filament and enhances nebulin’s vulnerability to proteolysis.^30,69^ Hence, normal transcript but reduced protein levels in some of NEM2 patients might be due to such structural mismatch between nebulin and the thin filament. This can explain the reduced nebulin levels in patients with full exon deletion which leads to partial SR deletion. For instance, patient 144, who is homozygous for the deletion of exon 55, has normal transcript levels but reduced protein levels which is likely caused by partial deletion of SR9. Similarly, nebulin levels might be reduced in patient 151 (82% skipping of exon 30) due to partial deletion of SR2. Disruption of binding sites on nebulin can also be caused by activation of cryptic splice sites and subsequent intronic inclusions. For instance, in patient 4001, the *NEB* transcript appears normal, yet the protein level is diminished. Prediction of nebulin structure in this patient revealed a partial intronic inclusion at the site of the splicing mutation and this results in an unstructured region which disrupts the actin binding sites on nebulin (Fig. 2B).

Splicing analysis of patient 2385 revealed that transcripts in this patient have either exon 77 skipped, or exon 32 skipped, or they have cryptic splice site usage that results in an out-of-frame transcript due to a 7 bp deletion (Fig. 2A and Supplementary Table 2). It is to be expected therefore that due to the above discussed nebulin-thin filament structural mismatch, nebulin protein is degraded, yet protein levels are normal in this patient (Fig. 3B). We speculate that considering that skipping exon 32, which encodes SR3 which is close to the nebulin’s end, may not be as harmful as skipping exons located more centrally. Another potential scenario could be that post-translational modifications may contribute to the incorporation of mutated nebulin into thin filaments, rendering it resistant to protein degradation. Further investigation is required to explain the normal nebulin level in this patient.

Mutations affecting donor splice sites were identified in five NEM2 patients (3424, 2622, 4001, 151 and 2385). The transcripts originating from these alleles exhibited partial intron inclusion and the activation of cryptic splice sites (Fig. 2A and Supplementary Table 2). Given that splice site mutations are the most prevalent type of mutation within the *NEB* gene,^70^ it is important to consider the potential impact of these frequently occurring cryptic splice sites when interpreting identified *NEB* gene variations. In our cohort, two intronic variants were initially classified as VUS. However, our results demonstrate that both variants have pathogenic implications, either by inducing exon skipping (patient 151) or by activating cryptic splice sites and intron inclusion (patient 3424). These findings underscore the need to optimize cryptic splice-site prediction algorithms or perform transcriptomic studies for a more accurate interpretation of the functional implications of splice site and intronic mutations within the *NEB* gene.

Our findings revealed allelic imbalance for all truncation mutations in our patient’s cohort (Fig. 1B), showing their deleterious impact on *NEB* mRNA stability and their role in triggering NMD. This conclusion was further substantiated through protein analysis, as we were unable to detect any truncated nebulin protein by western blot (Fig. 3C). Absence of truncated proteins is consistent with the notion that transcripts containing premature stop codons (PTC) undergo degradation via the NMD pathway and support the role of NMD in regulating *NEB* transcripts. It is known that not all PTCs trigger NMD ^71^ and that the position of PTC,^72,73^ gene, length of exons carrying PTC mutation^74,75^ and tissue type^76,77^, all determine the efficiency of NMD. As an example, several studies have shown that *TTN* truncating variants (TTNtv) are not subject to substantial NMD^78-80^ which implies inefficiency of NMD to degrade TTNtv transcripts^81^ and as a result truncated proteins can be produced^78,79,82-84^. Furthermore, the allelic ratio for truncation mutations in our study is not zero (0.15-0.35) which suggests that the absence of truncated nebulin is also likely to include more effective protein degradation of truncated nebulin compared to truncated titin.

### Effect of *NEB* mutations on TFL and tension production

Nebulin’s role in TFL regulation and force generation is well studied in mouse models.^26,30,31,34,35,37,26-31^ Here we addressed how nebulin mutations influence TFL and force in NEM2 patients. Results underscore the pivotal role of nebulin in governing TFL, as evidenced by a notable negative relation between loss of nebulin and TFL (Fig. 4C). Patients with reduced nebulin levels exhibited significantly shorter TFL in comparison to controls. Conversely, patients with normal nebulin demonstrated TFL similar to controls (Fig. 4B). Furthermore, patient 180, with a larger nebulin size, exhibited a longer TFL relative to controls (Fig. 4D). RNA-seq results showed transcripts of this patient with 2859 additional bp comparing to controls which equates to two more TRI copies (Supplementary Table 3), or four additional SRs. TFL can be expected therefore to be longer than in controls, and this is what was measured (Fig. 4D-right). Longer TFL in this NEM2 patient reveals, for the first time, that not only shortened but lengthened TFL is implicated in the pathomechanism of NEM2. This finding aligns with previous study which suggested that *NEB* can tolerate deviations of one TRI copy, whereas the addition of multiple copies may be pathogenic.^50^ Our studies also showed that the transcript level of this patient is reduced (Fig. 1C), but nebulin protein level is relatively normal (Fig. 3B). It is possible that the two-copy gain may disrupt the stability or secondary structure of the mRNA,^50^ which is reflected in the lower level of *NEB* transcript, but does not impact protein level, possibly due to tighter binding between longer nebulin and thin filament which increases protein stability.

We observed a negative relation between reduction of nebulin level and tension production, both for maximal and submaximal tensions (Fig. 5). This correlation underscores that patients with relatively normal nebulin exhibit a less pronounced reduction in tension compared to those with reduced nebulin. The diminished tension observed in patients with nebulin deficiency can be attributed to factors such as altered cross-bridge cycling kinetics, changes in thin filament and sarcomere structure and myofibrillar misalignment.^26-31^ Our finding that force is reduced in some patients despite a relatively normal level of nebulin bears resemblance to the findings in a mouse model of typical nemaline myopathy^85^ wherein a decrease in specific force was noted despite the presence of normal nebulin levels. The force deficit observed in this mouse model was attributed to changes in thin filament structure and less organized myofibrils^85^ and a similar explanation might hold here.

It is also relevant to note that TFL was determined in the present study by using optical techniques, and this provides an average length but cannot determine the variation in thin filament length. In previous immunoelectron microscopy studies on mouse muscle it was shown that when nebulin levels are reduced, thin filaments are shorter *and* vary greatly in length. The shorter length and length variation lowers the force on the descending limb of the force-sarcomere length relation as well as lowers the maximal force at optimal sarcomere length (see Fig. 3 in^27^). Thus, it is likely that both the reduction and variation in TFL explains the effects on tension that we measured, with the most severe effects in patients with the lowest level of nebulin because the TFL is most severely affected. Patient 180 is unique because it has thin filaments that are *longer* than normal. The consequence of longer TFL is that the force-sarcomere length relation is right shifted^86^ and that the operating sarcomere length might include the ascending limb where force is less than optimal.

In summary, the studies on NEM2 patients support the critical importance of a normal level of full-length nebulin for thin-filament length regulation and force production. Both a reduction and an increase in TFL are deleterious, and the uniformity in TFL is critical as well.

### Effect of OM on force production

OM is a small-molecule activator that was developed for the treatment of heart failure which increases the calcium sensitivity of force production by binding to cardiac myosin (*MYH7*).^44,45,59^ OM is also effective on type I skeletal muscles,^46^ because these fibers express the same myosin isoform as found in cardiac myocytes.^87^ Given the large number of type I fibers in humans,^88,89^ and the additional shift in NEM patients toward type I fibers,^4,5^ OM is an attractive candidate to counteract muscle weakness in NEM2 patients.

OM treatment increases calcium sensitivity (Fig. 6A and B) and consequently submaximal force production (Fig. 6D) of slow fibers in all NEM2 patients. This finding is consistent with the mechanism proposed for OM which suggests OM binds to myosin, leading to long-lasting, inactive myosin heads that cooperatively activate the thin filaments and trigger binding of additional myosin heads that increase submaximal force.^90^ The dynamic stiffness measurements in OM treated fibers were aligned with this mechanism revealing significantly higher ED of OM-treated slow fibers of both patients at pCa 6.75 relative to vehicle and no change of ED at pCa 4, indicating that at submaximal level of activation and in presence of OM, there are more strongly bound cross bridges. Moreover, it has been shown that OM disrupts the typical pathway of actin-myosin interaction, causing the detachment of myosin heads independent of ATP hydrolysis.^90^ Thus, it can be postulated that OM treatment leads to reduction of tension cost due to long-lasting cross-bridges and ATP-independent detachment of cross-bridges. Our results of dynamic stiffness measurement agree with this notion as they show in OM-treated fibers slower *c*, supporting the effect of OM in reducing tension costs (Fig. 7C and Supplementary Fig. 6A).

Interestingly, OM enhanced submaximal force generation to a higher degree in NEM2 patients than in control fibers (Fig. 6F). This finding aligns with the study of Lindqvist et al^46^ in which OM had a greater effect in slow fibers from Neb cKO mice than from control fibers. The correlation between tension enhancement following OM treatment and the nebulin deficit may be explained by the different effects of nebulin mutations on *c*. Our results show that OM lowers *c* in the studied patient with reduced nebulin to a greater extent compared to controls or patient with normal nebulin (Supplementary Fig. 7). A lower *c* value indicates longer lasting myosin cross-bridges in the strongly bound state and thereby more force production. Whatever the underlying mechanism, the effect should be beneficial in patients with the lowest nebulin level who have the most severe force reduction and having a drug that has a great effect in these types of patients is highly needed.

For OM to be a beneficial therapy in NEM2 patients, it is important to avoid adverse effects. For instance, our findings demonstrate that 0.5 µM OM treatment reduces *k*_tr_ and *c* (Fig. 7A and C). Such reduction in cross-bridge cycling kinetics will adversely affect relaxation, which can be harmful in the heart. However, clinical trials have shown that OM is well tolerated without adverse effects up to ∼1 μM of plasma concentrations in heart failure patients.^91^ Thus, it is possible that the effects established in the present study using 0.5 µM OM will be able to increase skeletal muscle force in NEM2 patients without adverse cardiac effects.

In summary, NMD plays a crucial role in regulating *NEB* transcripts in patients with truncation mutations, and a high incidence of cryptic splice site activation and intronic inclusion was found in patients with splicing mutations. Intronic inclusion was shown to lead to insertion of an unstructured sequence into actin binding motifs and we propose that this disrupts the proper domain spacing and negatively impacts nebulin function. Considering that nebulin consists of a long chain of repeating units, each with thin filament binding sites that need to match with the regularly spaced thin filament proteins, the local displacement of a binding site is expected to have long range effects. Thus, a protein like nebulin might have heightened sensitivity to cryptic splice site activation. Additionally, our study underscored the importance of proper TFL, and that both shorter and longer than normal length can be detrimental. Finally, treatment with OM substantially increased force production in NEM2 patients, and the effect is largest for patients with the lowest level of nebulin. Given the absence of a curative treatment for NEM2, these results provide a foundation for future investigations into the potential therapeutic benefits of OM for NEM patients.

## Data availability

Raw data were generated at the University of Arizona. Derived data supporting the findings of this study are available from the corresponding author on request.

## Acknowledgements

We are grateful to Shawn Granzier-Nakajima for developing dynamic stiffness software.

## Supplementary material

Supplementary material is available at *Brain* online.

