## Supplementary Tables, Figures and Materials and Methods for "Characterization of *NEB* mutations in patients reveals novel nemaline myopathy disease mechanisms and omecamtiv mecarbil force effects"

### Supplementary Figures

A

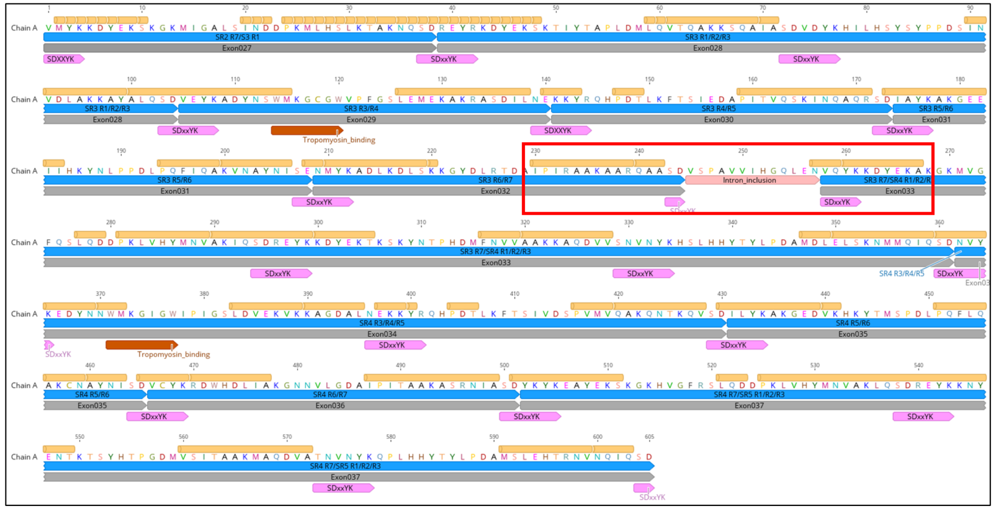

B

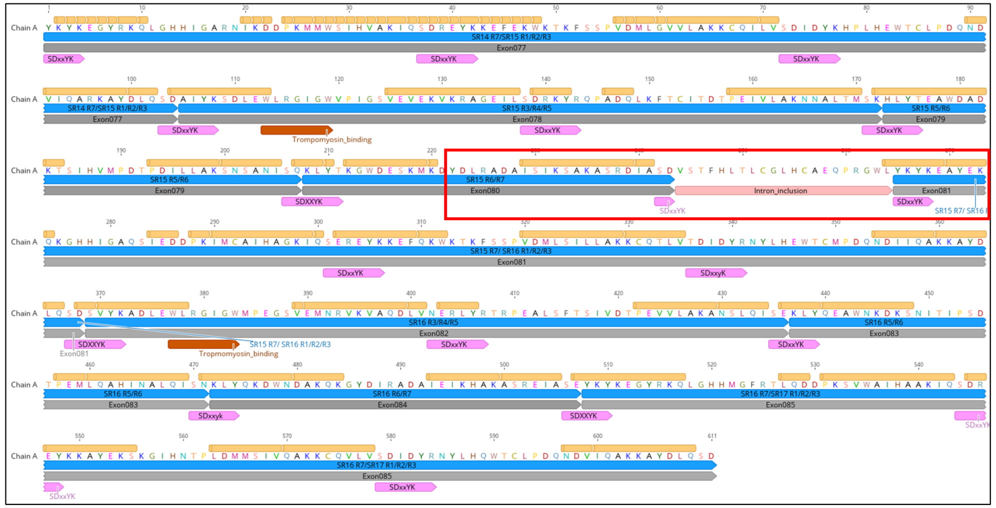

**Supplementary Figure 1. Predicted structure of mutated nebulin in patients 4001 and 2622.** The regular arrangement of nebulin's simple repeats, which include actin binding sites, is interrupted by the partial inclusion of intron 32 in patient 4001 (A) and intron 80 in patient 2622 (B). Predicted nebulin α-helix structure is in yellow, super-repeat region is in blue, acting-binding motif is in pink and exon is in grey.

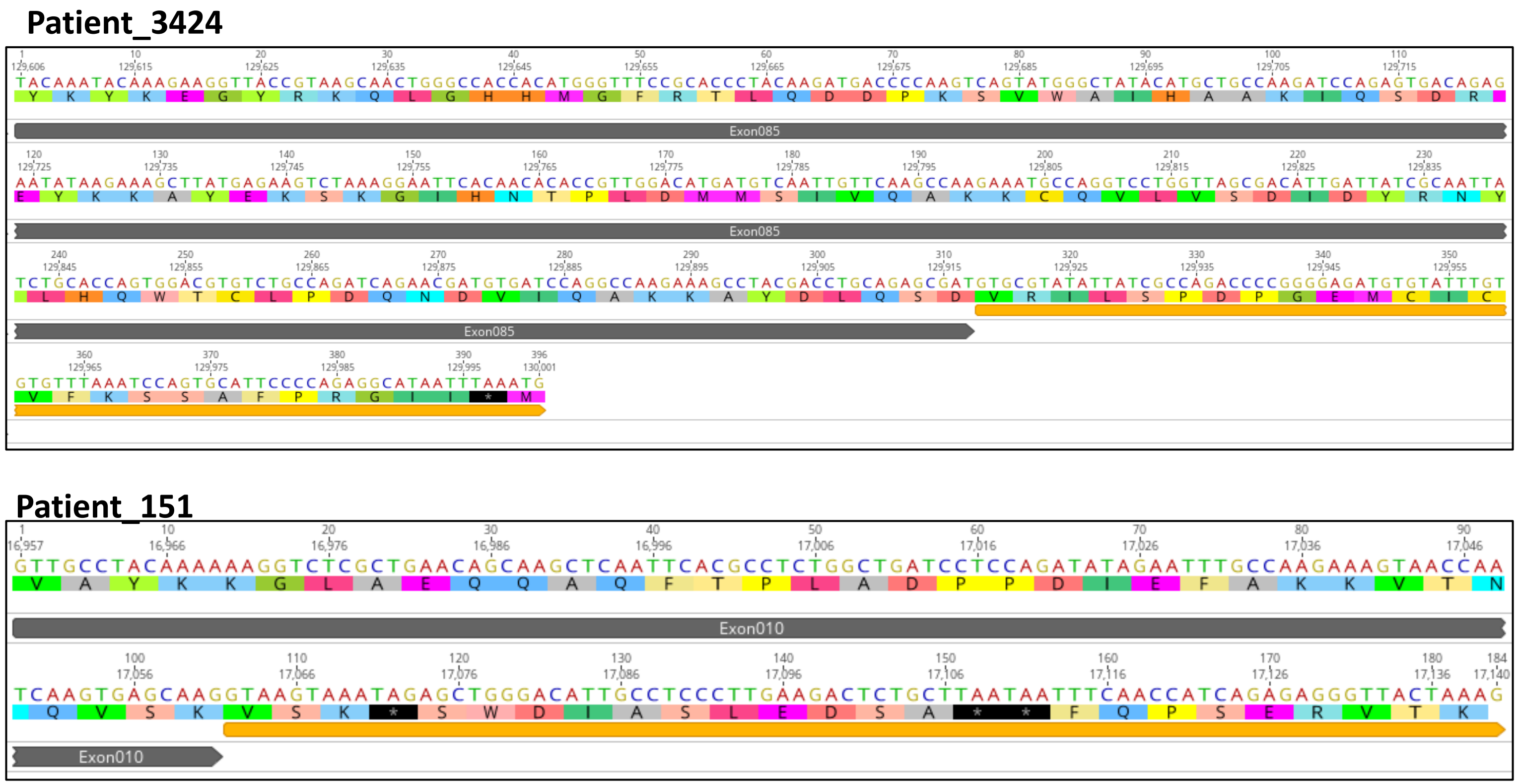


**Supplementary Figure 2. Intron inclusion outcome in patients 3424 and 151.** The top panel illustrates the in-frame partial inclusion of intron 85 in patient 3424, while the bottom panel shows the out-of-frame partial inclusion of intron 10, both triggering a premature stop codon and subsequent transcript degradation through NMD.

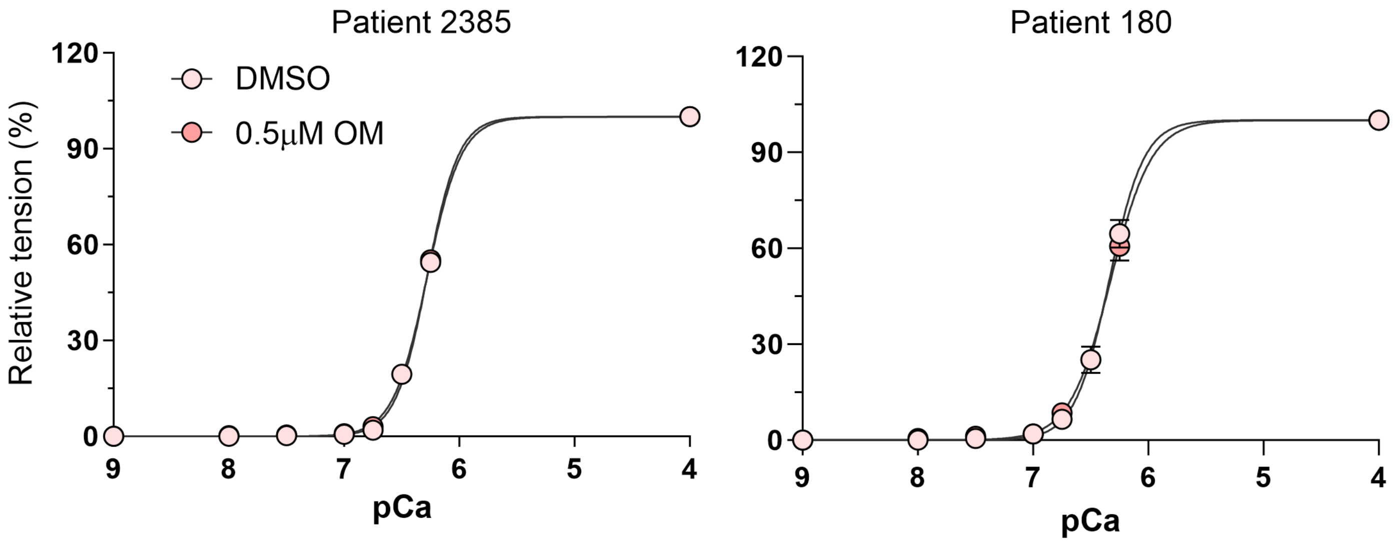


**Supplementary Figure 3. Effect of Omecamtiv mercarbil (OM) on type II fibers from NEM2 patients.** 0.5 μM OM treatment of fast fibers (Type II) does not affect the force-pCa relation.

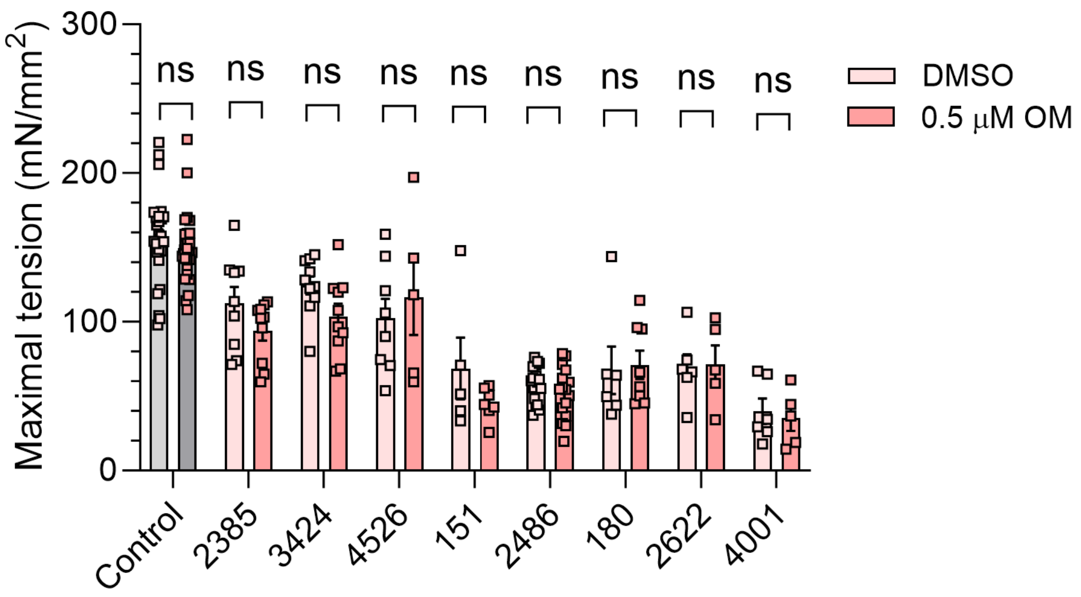


**Supplementary Figure 4. Effect of Omecamtiv mercarbil (OM) on maximal tension (pCa 4).** The maximal tension of controls and NEM2 patients (regardless of nebulin level) is not changed by 0.5 μM OM treatment. ns indicates no significant difference.

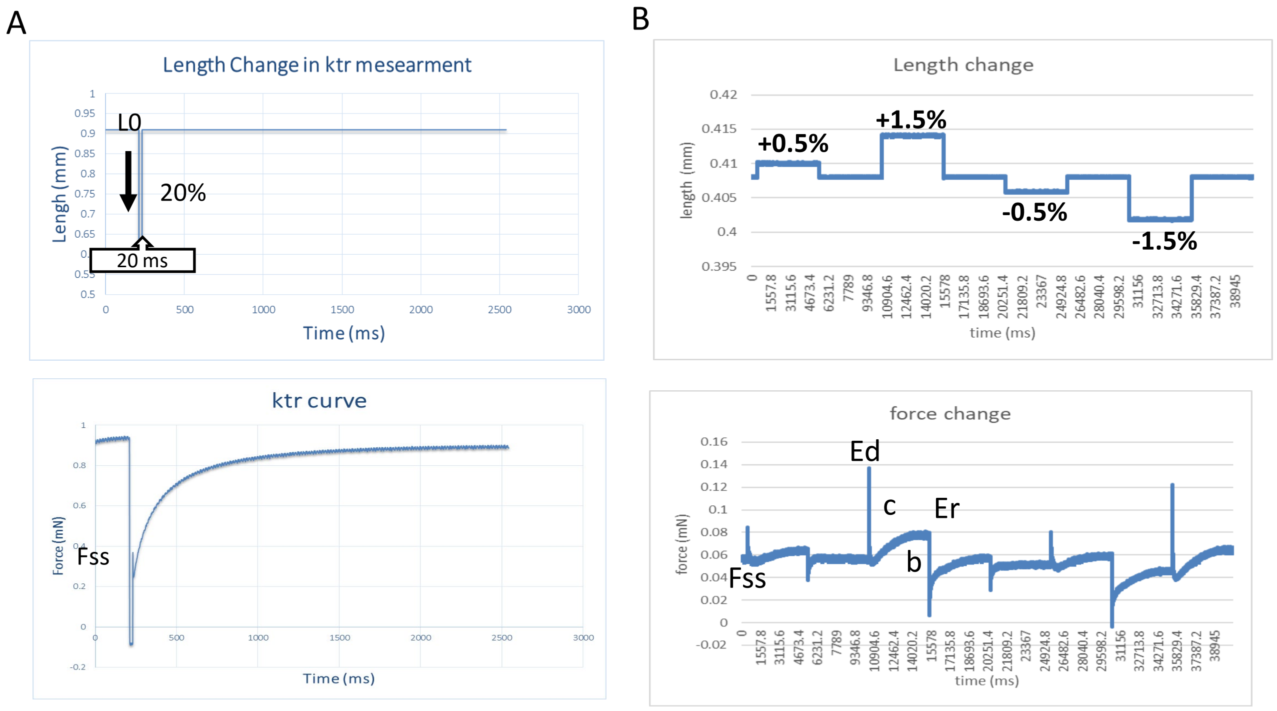


**Supplementary Figure 5. Protocol and tension transient for rate of tension redevelopment (*ktr*) and dynamic stiffness measurements.** A) Fiber length change (top) and force trace (bottom) for measuring ktr. At steady-state force (Fss) the muscle fiber was rapidly (<1 ms) shortened by 20% at one end of the fiber. After 20 ms the fiber was stretched to its initial length. ktr was determined by fitting the rise of force to the following equation: $F = Fss*(1-ektr*t)+c$, where F is force at time t, Fss is steady-state force, B) Analysis of Dynamic Stiffness: At Fss, a sequence of rapid release and stretch perturbations was introduced to the fiber preparations (Top). Subsequently, the distinct phases of tension transients (Bottom) that emerged due to alterations in muscle length were individually examined to gain insights into cross-bridge dynamics. The force changes were analyzed using a non-linear recruitment-distortion (NLRD) model, yielding values for Ed (an approximation of strongly bound cross-bridges), Er (an approximation of newly formed cross-bridges), *c* (an approximation of detachment rate), and *b* (an approximation of cross-bridge attachment rate).

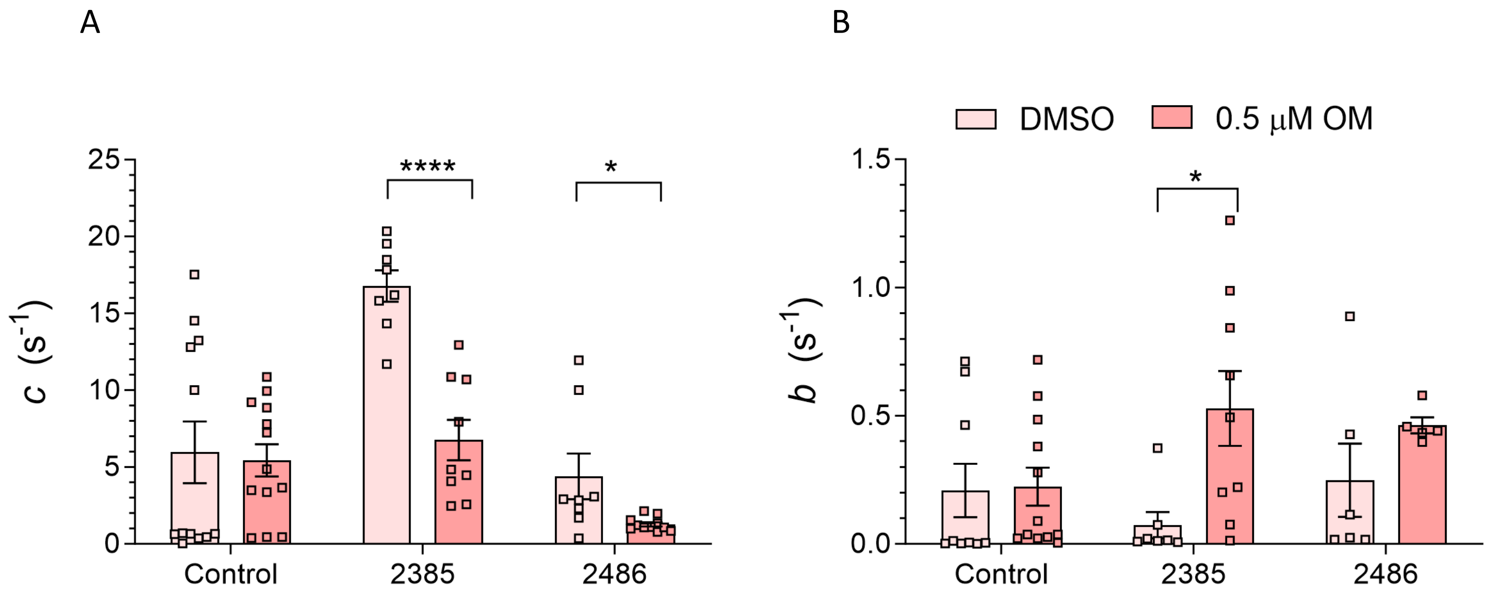


**Supplementary Figure 6. Effect of Omecamtiv mercarbil (OM) on dynamic stiffness at submaximal level of activation (pCa6.75).** A) OM treatment lowers the detachment rate of cross-bridges (c) in patients but not controls. B) OM treatment increases attachment rate of cross-bridges (b) in patient 2385. Asterisks indicate a significant difference between DMSO and OM-treated fibers and ns indicates no significant difference. * P <0.05, ** P <0.01, *** P <0.001, **** P <0.0001.

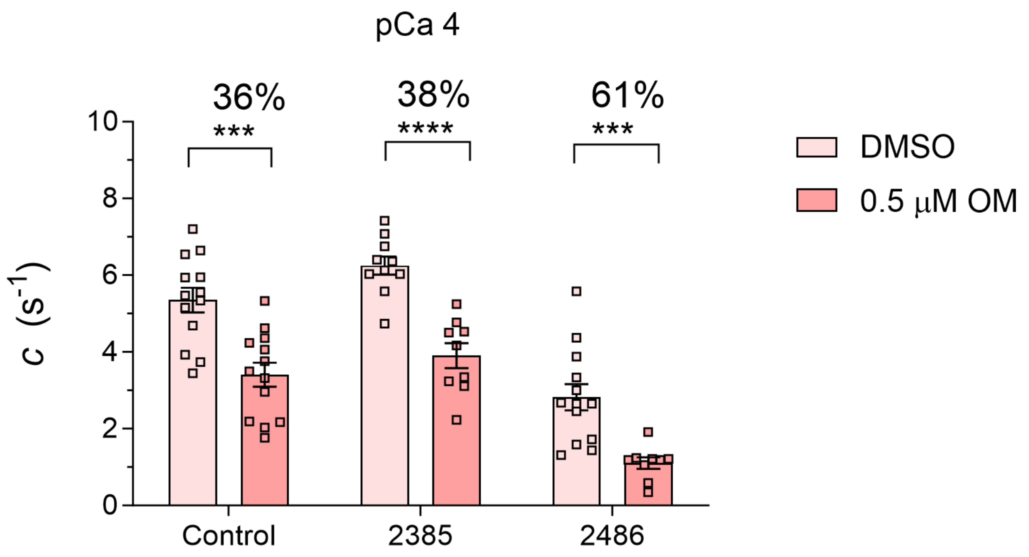


**Supplementary Figure 7. Nebulin level-dependence of OM effect on Cross-Bridge Detachment Rate (*c*).** OM-treatment lowers c in the patient with less nebulin (2486) to a greater extent (61%) than the patient with Normal nebulin (2385) or control (38 and 36% respectively).

### Full blots of Figure 3 C

N terminal

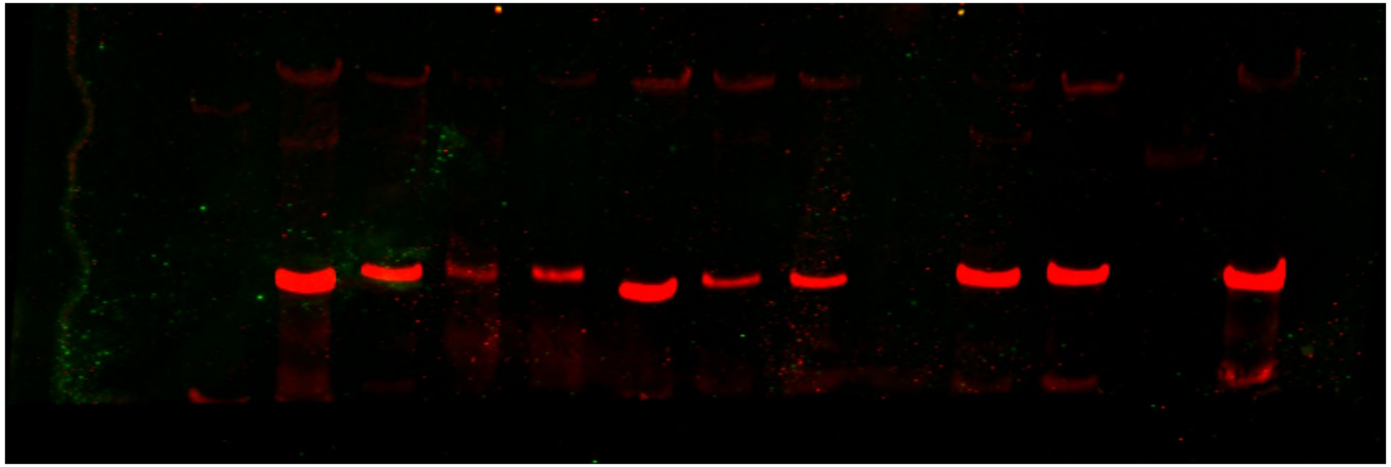

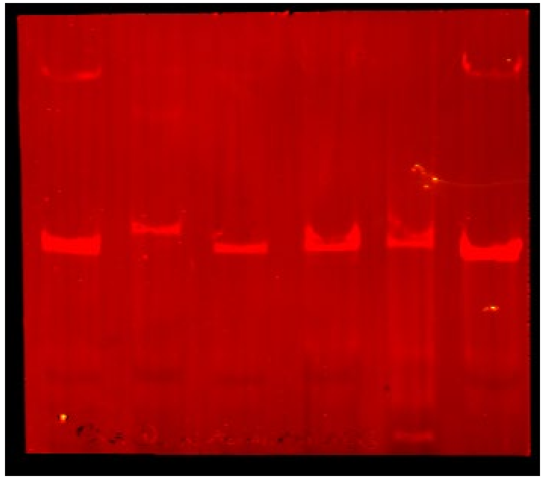

C terminal

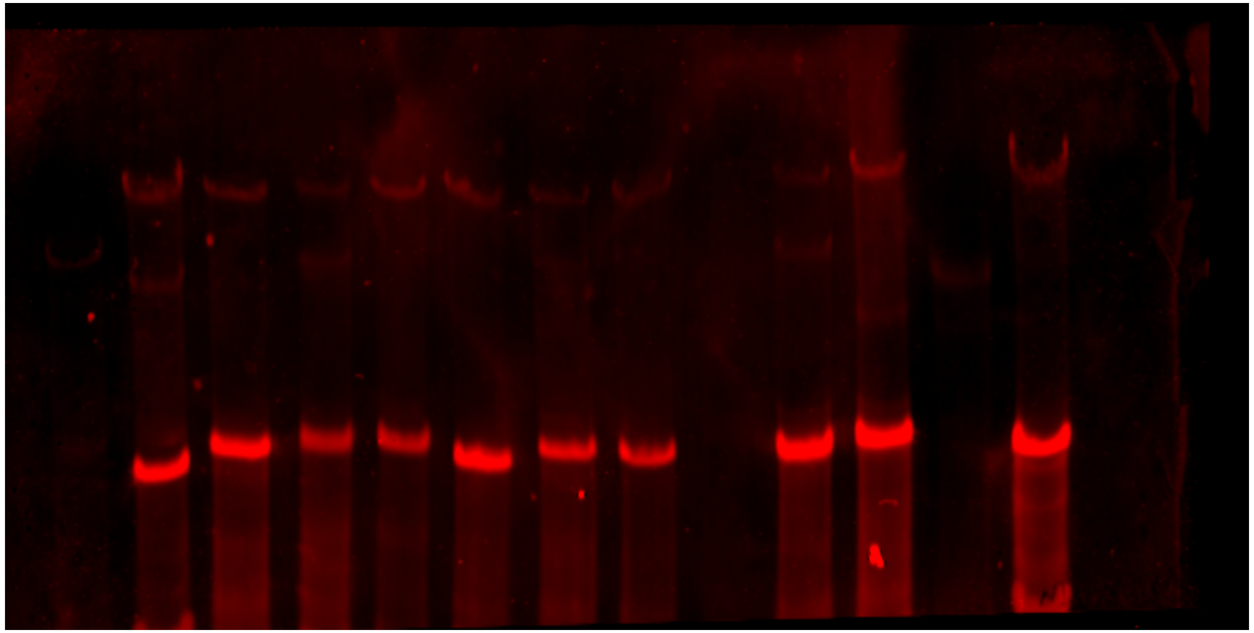

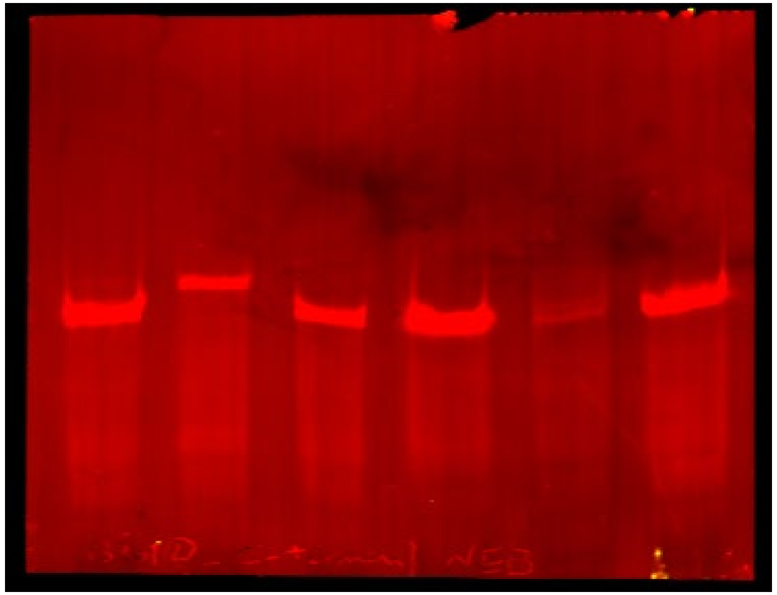

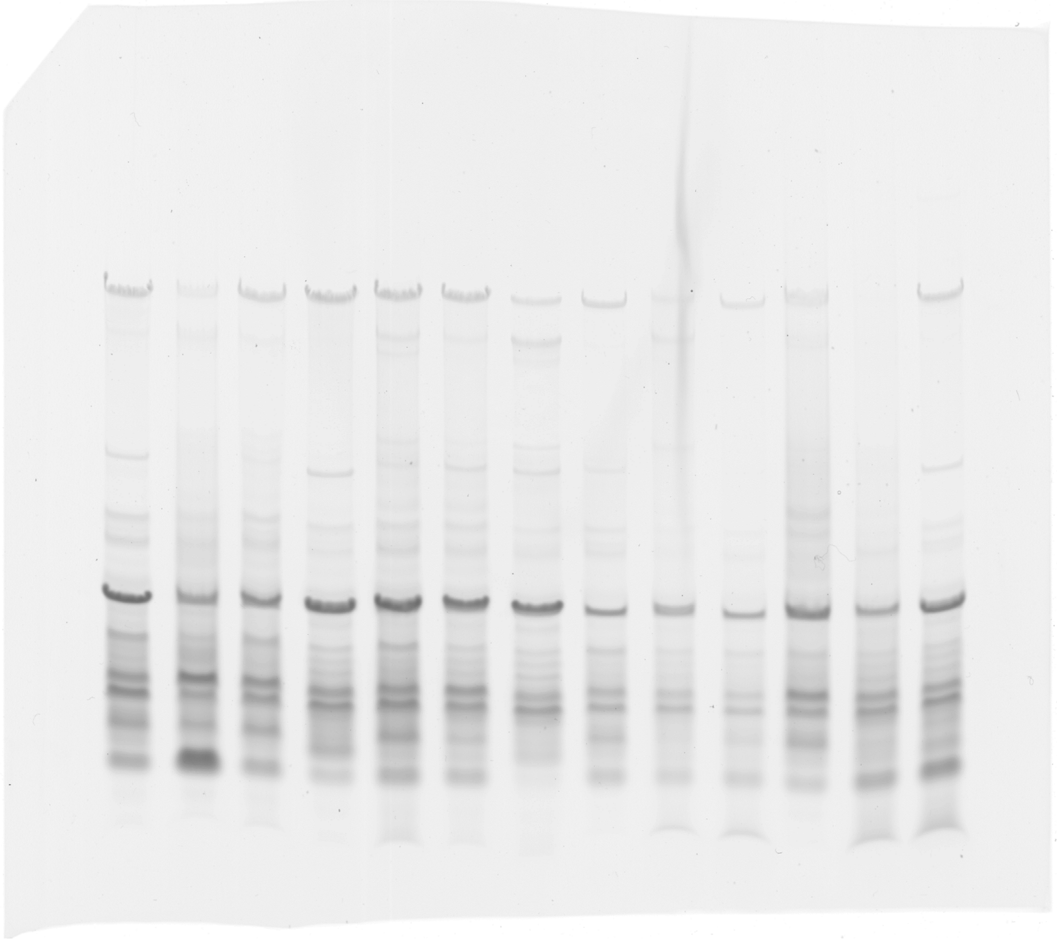

**Full gel for Figure 3 A**

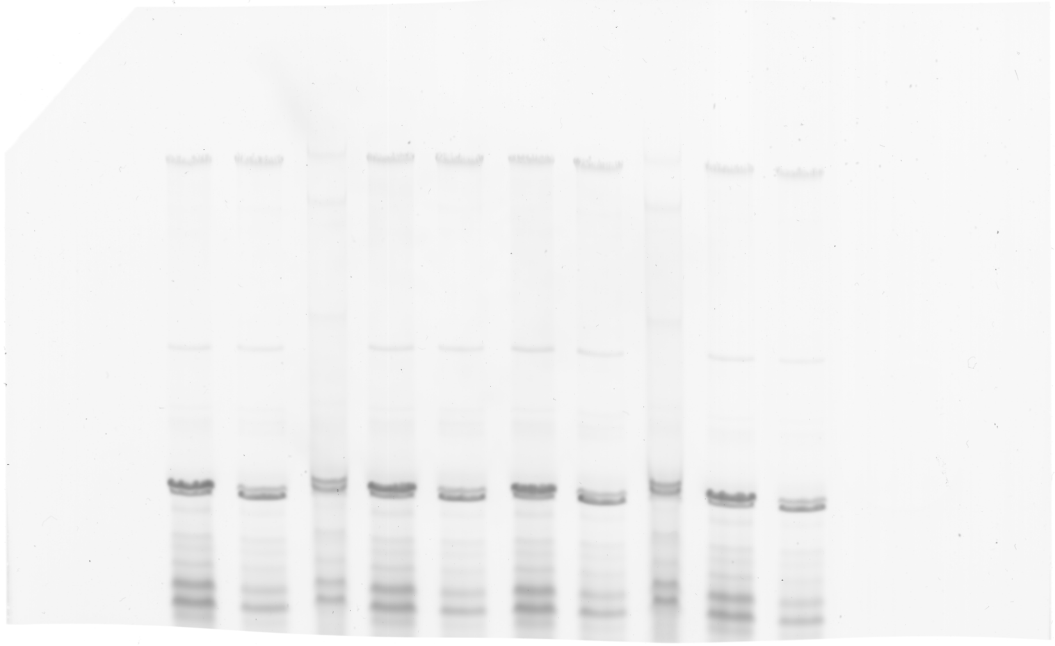

**Full gel for Figure 4 D left**

**Supplementary Table 1. Clinical characteristics of patients with NEM2**

| Patient ID | Site of biopsy | Age (yrs) at biopsy | Gender | Jump both feet leaving ground | Walk unassisted | Sitting unassisted | Breathing tube and vent in neonatal period | Feeding difficulties | Assisted breathing |
|---|---|---|---|---|---|---|---|---|---|
| 2385 | Paraspinal | 22 | F | Never able | Yes, greater than 10 steps | Unassisted more than an hour | No | Has or had G-tube & poor weight gain | Yes, > 12 hours/day |
| 2296 | paraspinal | 11 | F | Never able | Never able | Unassisted more than an hour | No | Has or had poor weight gain | Yes, > 12 hours/day |
| 2486 | Paraspinal | 5.75 | F | Never able | Never able | Unassisted more than an hour | No | Has or had G-tube & poor weight gain | Yes, > 12 hours/day |
| 3424 | right thigh | 7 | F | Yes | Yes, greater than 10 steps | Unassisted more than an hour | No | None | Yes, < 12 hours/day |
| 4001 | right thigh | 0.3 | M | Never able | Never able | Unassisted more than an hour | No | Has or had G-tube & poor weight gain | Yes, > 12 hours/day |
| 2622 | quadriceps | 0.75 | F | Never able | Never able | Unassisted more than an hour | No | Has or had NG tube | Yes, < 12 hours/day |
| 4526 | Left thigh | 2 | M | Never able | Never able | Unassisted more than an hour | No | Has or had G-tube & poor weight gain | Yes, < 12 hours/day |
| 144.0 | left tricep | 1.3 | F | Never able | Never able | Unassisted more than an hour | No | Has or had G-tube & poor weight gain | Yes, < 12 hours/day |
| 180.0 | left quad | 0.66 | F | Never able | Yes, greater than 10 steps | Unassisted more than an hour | No | Has or had G-tube & poor weight gain | Yes, > 12 hours/day |
| 151.0 | left thigh | 1 | F | Never able | Never able | Unassisted more than an hour | No | Has or had G-tube & poor weight gain | None |

**Supplementary Table 2. Summary of mutation analysis in NEM2 patients**

| Patient | Mutation | Mutation Type | Isoforms of transcripts | Transcript (%) in patients | Transcript (%) in controls | Cryptic splice site activation | Allelic imbalance | Intronic inclusion | Mutation site | PSI (%) | Clinical significance (ClinVar) | Total *NEB* transcript | Protein level |
|---|---|---|---|---|---|---|---|---|---|---|---|---|---|
| 2385 Allele 1 | exon 32 c.3252_3255+3delTGACGTA: Results 7 bp deletion | Deletion | Exon 31 ···· Exon 33 | 17.02 | 0.01 | | | | SR3 R6,R7 | 82 | Likely pathogenic | Normal | Normal |
| | | | Exon 32 ▬ Exon 33 | 9.73 | 0.02 | Yes | | Out-frame inclusion of 39 bp from intron 32 equates to addition of 13 amino acids | | | | | |
| 2385 Allele 2 | 2.7 kb deletion exon 77 g.152,469,299 in intron 77 g.152,471,995 in intron 76: Results 2697 deletion including exon 77 | Deletion | Exon76 — Exon78 | 49.6 | 0.03 | | | | SR14 R7 / SR15 R1,R2,R3 | 46 | - | | |
| 2296 Allele 1 | exon 112 c.17654G>A: Results generation of stop codon in exon 112 | Truncation | ▼ Exon 112 | 27 | | | Yes | | SR24 R2/R3 | 100 | Pathogenic | Reduced | Reduced |
| 2296 Allele 2 | exon 175 c.24771delT: Results deletion in exon 175 leading to stop 19 bp into exon 176 | Frameshift | ▼ Exon 176 | 39.34 | | | | | M234/M235 | 100 | Pathogenic/ Likely pathogenic | | |
| 2486 Allele 1 | exon 61 c.8425 C>T : Results generation of stop codon in exon 61 | Truncation | ▼ Exon 61 | 34 | | | Yes | | SR10 R7 / SR11 R1,R2,R3 | 100 | Pathogenic/ Likely pathogenic in LOVD | Reduced | Reduced |
| 2486 Allele 2 | exon 171 c.24317 T>A: Results generation of stop codon in exon 171 | Truncation | ▼ Exon 171 | 35 | | | Yes | | M230/M231 | 77 | Pathogenic/ Likely pathogenic in LOVD | | |
| | | | Exon 167 ···· Exon 173 | 30.5 | 22.8 | | | | | | | | |

| Patient | Mutation | Mutation Type | Isoforms of transcripts | Transcript (%) in patients | Transcript (%) in controls | Cryptic splice site activation | Allelic imbalance | Intronic inclusion | Mutation site | PSI (%) | Clinical significance | Total NEB transcript | Protein level |
|---|---|---|---|---|---|---|---|---|---|---|---|---|---|
| 3424 Allele 1 | exon 85 c.13059+5G>A: Results point mutation in intron 85 | Intronic point mutation | 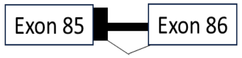 | 20.38 | 0.03 | Yes | | In-frame intron inclusion of 84 bp from intron 85 equates an addition of 28 amino acids. The inclusion would add a stop codon before exon 86. | SR16 R7 / SR17 R1,R2,R3 | 100 | Likely pathogenic | Reduced | Normal |
| 3424 Allele 2 | exon 169 c.24218C>A: Results generation of stop codon in last codon of exon 169 | Truncation | 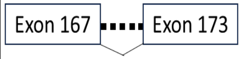 | 74.5 | 44.28 | | Yes | | M228/M229 (Z-disk) | 31 | VUS/ pathogenic in LOVD | | |
| | | | 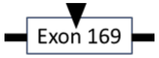 | 25.42 | 55.72 | | | | | | | | |
| 4001 Allele 1 | exon 32 c.3255+1G>A: Results donor splice mutation in junction of exon-intron 32 | Splicing | 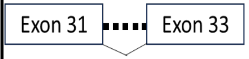 | 12.55 | 0.01 | | | | SR3 R6/R7 | 85 | Pathogenic | Normal | Reduced |
| | | | 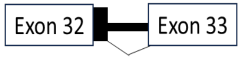 | 51.82 | 0.02 | Yes | | In-frame inclusion of 39 bp from intron 32 equates to an additional 13 amino acids | | | | | |
| 4001 Allele 2 | exon 110 c.17501_17502delinsC: Results insertion in exon 110 leading to stop codon 19 bp into exon 111 | Frameshift | 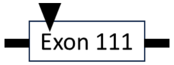 | 22.18 | | | | | SR23 R7/SR24 R1 | 100 | Pathogenic in LOVD | | |
| 4526 Allele 1 | exon 109 c.17262G>A:Results generation of stop codon in the middile of exon 109 | Truncation | 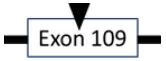 | 23.9 | | | Yes | | SR22 R6/R7 | 100 | Pathogenic | Reduced | Normal |
| 4526 Allele 2 | exon 171 c.24318_24319insAA: Results insertion at the start of exon 171 leading to stop codon in exon 173 | Frameshift | 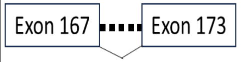 | 73.87 | 22.8 | | | | M229/M230 (Z-disk) | 53 | Pathogenic | | |
| | | | 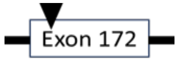 | 40 | | | | | | | | | |

| Patient | Mutation | Mutation Type | Isoforms of transcripts ** | Transcript (%) in patients | Transcript (%) in control | Cryptic splice site activation | Allelic imbalance | Intronic inclusion | Mutation site | PSI (%)* | Clinical significance | Total NEB transcript | Protein level |
|---|---|---|---|---|---|---|---|---|---|---|---|---|---|
| 2622 Allele 1 | exon 80 c.12018+1G>A: Results donor splice site mutation in juncation of exon-intron 80 | Splicing | 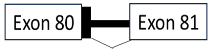  | 4.58 | 0.01 | Yes | | In-frame inclusion of 63 bp from intron 80 equates to 21 additional amino acids | SR15 R6/R7 | 80 | Pathogenic/Likely pathogenic | Reduced | Reduced |
| | | | 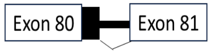  | 3.27 | 0 | Yes | | Out-frame inclusion of 95 bp from intron 80 | | | | | |
| 2622 Allele 2 | exon 172 c.24458_24461dupAGAT: Duplication of AGAT in exon 172 leads to stop codon at the end of exon | Frameshift | 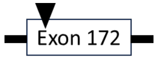  | 25 | | | | | M231/M232 | 100 | Pathogenic/Likely pathogenic | | |
| 144 Allele 1 & 2 | Homozygous exon 55 deletion: Results in-frame 2502 deletion including exon 55 | Deletion | 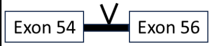  | 100 | 0 | | | | SR9 R5,R6 | 4 | Pathogenic | Normal | Reduced |
| 180 Allele 2 | Four copy gain within the *NEB* gene triplicate repeat region | Duplication | 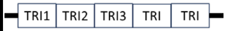  | 100 | 0 | | | | [SR16 R3,R4,R5] – [SR21 R7 / SR22 R1,R2,R3] | - | Pathogenic | Reduced | Normal |
| 180 Allele 1 | exon 157 c.22936C>T (p.Arg7646*): Results generation of stop codon in exon 157 | Truncation | 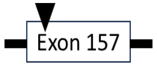  | 17 | | | Yes | | M216/M217 | 100 | Pathogenic | | |
| 151 Allele 1 | exon 10 c.822+1G>A: Results donor splice site mutation at the junction of exon-intron 10 | Splicing | 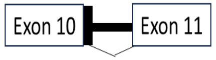  | 13.66 | 0.02 | Yes | | Out-frame inclusion of 79 bp from intron 10. The inclusion would add 3 stop codons before exon 11. | M5 | 100 | Likely pathogenic | Reduced | Reduced |
| 151 Allele 2 | exon 30 c.3042+3_3042+6del : Results 4 bp deletion of intron 30 | Intronic Deletion | 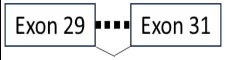  | 82.05 | 0.09 | | | | R5 of SR2 | 15 | VUS | | |

*PSI represents the inclusion percentage of the exons carrying mutations.

**Transcript isoforms are only showing isoforms are impacted by mutation and isoforms encoded by exons without mutations have not been shown.

Intron ▬, Stop codon ▼, Exon skipping ▪▪▪▪▪▪▪▪, Intron inclusion ┣━, Deleted exon V

#### Supplementary Table 3- Calculation of length of triplicated region in patient 180

| Exon NO | J1 | J2 | Reads per Exon (J1+J2) | Exon length | Normalized read count (J1+J2)/exon length | Normalized read count in controls | Ratio of normalized read counts of patient/controls | Exon length based on additional reads (ration of normalized counts * exon length) | Added exon length in patient (exon length based on additional counts - exon length) |
|---|---|---|---|---|---|---|---|---|---|
| Exon082 | 60305 | 7548 | 67853 | 203 | 334.2512315 | 201.53038 | 1.658565003 | 336.6887 | 133.6886957 |
| Exon083 | 14857 | 60279 | 75136 | 104 | 722.4615385 | 383.10897 | 1.885785995 | 196.1217 | 92.1217435 |
| Exon084 | 38236 | 14669 | 52905 | 107 | 494.4392523 | 247.62928 | 1.996691366 | 213.646 | 106.6459762 |
| Exon085 | 36004 | 38242 | 74246 | 311 | 238.733119 | 133.44051 | 1.789060241 | 556.3977 | 245.3977349 |
| Exon086 | 10953 | 35998 | 46951 | 203 | 231.2857143 | 186.94745 | 1.237169634 | 251.1454 | 48.1454357 |
| Exon087 | 9576 | 10902 | 20478 | 104 | 196.9038462 | 253.36218 | 0.777163531 | 80.82501 | -23.17499273 |
| Exon088 | 9582 | 9599 | 19181 | 107 | 179.2616822 | 203.19315 | 0.882223074 | 94.39787 | -12.60213108 |
| Exon089 | 43509 | 9594 | 53103 | 311 | 170.7491961 | 105.41372 | 1.61980051 | 503.758 | 192.7579587 |
| Exon090 | 73903 | 43455 | 117358 | 203 | 578.1182266 | 309.8046 | 1.866073747 | 378.813 | 175.8129707 |
| Exon091 | 63138 | 74076 | 137214 | 104 | 1319.365385 | 744.16987 | 1.77293577 | 184.3853 | 80.38532007 |
| Exon092 | 57637 | 63030 | 120667 | 107 | 1127.728972 | 624.16511 | 1.806779898 | 193.3254 | 86.32544907 |
| Exon093 | 17387 | 57609 | 74996 | 311 | 241.1446945 | 137.51554 | 1.753581394 | 545.3638 | 234.3638135 |
| Exon094 | 41034 | 17400 | 58434 | 203 | 287.8522167 | 123.3908 | 2.332849824 | 473.5685 | 270.5685142 |
| Exon095 | 34050 | 40999 | 75049 | 104 | 721.625 | 259.23077 | 2.783716617 | 289.5065 | 185.5065282 |
| Exon096 | 36842 | 34099 | 70941 | 107 | 663 | 277.89097 | 2.385827831 | 255.2836 | 148.2835779 |
| Exon097 | 46125 | 36846 | 82971 | 311 | 266.7877814 | 121.88746 | 2.188804179 | 680.7181 | 369.7180996 |
| Exon098 | 70513 | 46128 | 116641 | 203 | 574.5862069 | 302.33333 | 1.900505646 | 385.8026 | 182.8026461 |
| Exon099 | 66109 | 70612 | 136721 | 104 | 1314.625 | 742.92628 | 1.769522807 | 184.0304 | 80.03037193 |
| Exon100 | 58142 | 66109 | 124251 | 107 | 1161.224299 | 647.68536 | 1.792883357 | 191.8385 | 84.83851915 |
| Exon101 | 16891 | 58142 | 75033 | 311 | 241.2636656 | 136.37406 | 1.769131621 | 550.1999 | 239.199934 |
| Exon102 | 12405 | 16908 | 29313 | 203 | 144.3990148 | 133.31034 | 1.083179366 | 219.8854 | 16.88541128 |
| Exon103 | 9821 | 12405 | 22226 | 104 | 213.7115385 | 252.02885 | 0.847964595 | 88.18832 | -15.81168212 |
| Exon104 | 9710 | 9821 | 19531 | 107 | 182.5327103 | 196.08411 | 0.930889853 | 99.60521 | -7.394785759 |
| Exon105 | 8004 | 9725 | 17729 | 311 | 57.00643087 | 69.453376 | 0.820787037 | 255.2648 | -55.73523148 |
| **Additional exon based in patient 180 versus control by summing up all added exon length in patients** | | | | | | | | | **2858.76** |

### Materials and methods:

#### Skeletal muscle biopsies of NEM2 patients

Surgically obtained muscle biopsy samples of NEM2 patients, from thigh, triceps or paraspinal were obtained from Congenital Muscle Disease Tissue Repository (CMD-TR) housed at the Medical College of Wisconsin, along with limited data including the mutations related to the specimens. We also obtained control biopsy samples from vastus lateralis muscles of 34 years old healthy individuals from the University of Montana.[1] The subjects' consent was obtained according to the Declaration of Helsinki and ethical approval for study of human biopsies was granted by Institutional Review Board of University of Arizona. All biopsies were stored frozen and unfixed at -80°C until use.

#### RNAseq analysis

Around 15-150 mg mg (depending on quality of biopsy) of frozen muscle tissue from NEM2 or control biopsy samples were collected and stored in RNAlator to reserve RNA integrity. For RNA extraction, 600 µl of prechilled buffer RLT (RNeasy Fibrous Tissue Mini Kit, Qiagen) with 1% β-mercaptoethanol was added to muscle tissue stored in RNAlater in a 4-ml cryovial. Tissue was disrupted using a rotor-stator homogenizer for 30 s. RNA extraction was performed following the manufacturer's instructions and quantified using a Nanodrop ND-1000 spectrophotometer (Thermo Fisher Scientific). RNA integrity was checked on a 2100 Bioanalyzer (Agilent), and all RNA integrity number scores were confirmed to be ≥8.

For library preparation, ribosomal RNA (rRNA) was depleted from RNA preparations with a NEBnext rRNA depletion kit using 1 µg of total RNA as starting material. Libraries were prepared using the NEBNext Ultra II Directional RNA Library Prep Kit for Illumina following the manufacturer's instructions. RNA was fragmented for 10 min at 94°C. For first strand cDNA synthesis, incubations were for 10 min at 25°C followed by 50 min at 42°C and 15 min at 70°C. For size selection, conditions for an approximate insert size of 300 bp were used. Size-selected libraries were enriched by PCR for 10 cycles and purified using NEBnext sample purification beads. Library quality and insert sizes were checked using a 2100 Bioanalyzer (Agilent). Sequencing was performed on an Illumina Hiseq2500 sequencer using 150-bp paired-end sequencing. The raw data are available (BioProject accession PRJNA996928). Adapters and low quality reads were removed with Trim Galore (www.bioinformatics.babraham.ac.uk/ projects/trimgalore/), and reads were mapped to the human genome (Release GRCh38.p13) using STAR[2] with default settings.

For calculating inclusion percentages of all exons from nebulin transcripts, inclusion reads (IRs) and exclusion reads (ERs) were counted for each exon based on nebulin isoform NM_001164508.2. IRs are reads overlapping the exon being investigated, normalized by exon length. ERs are reads either upstream or downstream that support exclusions of the read. From these factors, the following equations were used to calculate the PSI index using the ASpli R-package[3]

$IR_{i,n} = IR_i / length\ exon_i + read\ length - 1$
$ER_{i,n} = ER_i / read\ length - 1$
$PSI_i = (IR_{i,n} / IR_{i,n} + ER_{i,n})\%$

where *i* is the exon number and *n* is the normalized read counts.

Read density (normalized reads per base) was calculated by dividing read counts (per exon) by exon length. Read density was then divided by mean density in control samples. To estimate the number of added TRI repeats in patient 180, normalized read density was then multiplied by exon length and exon length was subtracted to estimate the number of additional bases per exon (Supplementary Table 3). The estimated number of added bases was then divided by the average repeat length to estimate the number of additional (super)-repeats.

For detection of alternative splicing events around mutations, the most frequent events were considered, and junction reads were used to calculate relative frequencies of events (exon skipping vs. intron inclusion vs. normal junction usage).

For allelic ratio calculations, base composition at single genomic positions was determined using the mpileup function from samtools.[4] Alleles at mutated positions were counted and ratios were calculated as follows: Allele 1 (mutated) / (Allele 1 (mutated) + Allele 2 (wt)). Following this calculation, equal expression from both alleles would give a ratio of 0.5. Degradation of the mutated allele by NMD would shift the ratio closer to zero and result in allelic imbalance.

#### Structure prediction with AlphaFold/Colabfold:

Alpha-helical portions of nebulin were predicted with Colabfold[5] using standard settings. As inputs for structure predictions, sequence lengths of two super-repeats around in-frame intron inclusions were selected (super-repeats 3-4 for patient 4001 and super-repeats 15-16 for patient 2622).

#### Omecamtive mecarbil

Omecamtiv mecarbil (OM) was purchased from Selleckchem (Houston, TX) and was dissolved in dimethylsulfoxide (DMSO) to make a stock solution as instructed by the manufacturer. OM stock solution was then added to the experimental solution to prepare the final desired concentration of OM. while keeping the concentration of DMSO at 0.5%. Based on our previous study that showed the force increase with OM reached a plateau between 0.5 and 1.0 µM, and with force starting to decrease at higher concentrations, we used 0.5 µM OM here.[6] 0.5% DMSO was used as vehicle.

#### Skeletal muscle mechanics

Small pieces cut from the frozen muscle biopsies were placed in 50% glycerol/relaxing solution (in mM: 40 BES, 10 EGTA, 6.56 MgCl2, 5.88 NaATP, 1 DTT, 46.35 K-propionate, 15 creatine phosphate, Ionic strength 180 mM, pH 7.0 at 20˚C) containing protease inhibitors (in mM: 0.01 E64, 0.04 leupeptin and 0.5 PMSF) and stored overnight at -20˚C. The solution was replaced with fresh 50% glycerol/relaxing solution the following day and the muscle tissue was membrane-permeabilized at 4˚C overnight as described previously.[7] This procedure renders the membranous structures in the muscle fibers permeable, which enables activation of the myofilaments with

exogenous calcium. Preparations were washed thoroughly with relaxing solution and stored in 50% glycerol/relaxing solution at -20°C. Muscles were used for experiments within 1 week. Small muscle bundles or single muscle fibers were dissected from the permeabilized strips and were mounted between a length motor (ASI 403A, Aurora Scientific Inc., Ontario, Canada) and a force transducer element (ASI 315C-I, Aurora Scientific Inc., Ontario, Canada) in a single fiber apparatus (ASI 802D, Aurora Scientific Inc., Ontario, Canada) that was mounted on the stage of an inverted microscope (Zeiss Axio Observer A1, Zeiss, Thornwood, NY, USA). Sarcomere length was set using a high-speed VSL camera and ASI 900B software (Aurora Scientific Inc., Ontario, Canada). Mechanical experiments were performed at a sarcomere length of 2.5 µm for all control and patients except patient 180 which we did mechanical experiments at sarcomere length of 2.7 µm. Fiber diameter and depth (determined with a build in prism that allowed for side view of the fiber) were measured at three points along the fiber, and the cross-sectional area was determined assuming an elliptical cross section. The temperature of the bathing solutions was kept constant at 20°C using a TEC controller (ASI 825A, Aurora Scientific Inc. Ontario, Canada). After completion of the mechanical studies (below), fibers/bundles were fiber-typed. Other than relaxing solutions, we used pre-activation (in mM: 40 BES, 1 EGTA, 6.32 MgCl2, 5.82 NaATP, 1 DTT, 81.71 K-propionate, 15 creatine phosphate, pH 7.0 at 20˚C) and activation solution (in mM: 40 BES, 10 CaCO3 EGTA, 6.29 MgCl2, 6.12 Na-ATP, 1 DTT, 45.3 potassium-propionate, 15 creatine phosphate, Ionic strength 180 mM, pH 7.0 at 20˚C) to do following mechanical experiments.
Force-pCa relationship: To determine the effect of 0.5 µM OM on the force-pCa relation, permeabilized muscle fiber bundles or single fibers were sequentially bathed in a relaxing solution, a pre-activation solution, and activation solutions with pCa values ranging from 8.0 to 4 - all containing 0.5 µM OM or vehicle - and the steady-state force was measured. The pCa-solutions were created by mixing relax and activating solutions taking into account the Kd of Ca2+ according to the model developed by Fabiato & Fabiato .[8] Measured force values were normalized to the maximal force obtained at pCa 4. The obtained force-pCa data were fit to the Hill equation (Y = 1/(1 + 10nH(pCa - pCa50))) where the pCa50 corresponds to the calcium concentration that yields half-maximal force and the Hill coefficient, nH, to myofilament cooperativity.[9]

*ktr*-measurements: The rate of tension redevelopment (*ktr*) was measured at steady-state force by rapidly shortening (1 ms) the fiber at one end of the fiber resulting in unloaded shortening of the fiber for 20 ms. Remaining bound cross-bridges were detached by rapidly restretching the fiber to initial length and the tension redeveloped [10]. ktr was determined by fitting the rise of force to the following equation (one-phase association curve): F = Fss*(1-ektr*t)+c, where F is force at time t, Fss is steady-state force [Supplementary Fig. 5A].

Step response protocol to measure dynamic stiffness: Muscle fibers were bathed in Ca2+ solutions (pCa6.75) with or without 0.5 µM OM treatment at 2.5 µm. Once the fiber preparations attained steady-state force, a series of rapid stretch and release length perturbations was applied [Supplementary Fig. 5B-Top] and then the various phases of the tension in response to muscle length (ML) changes [Supplementary Fig. 5B-bottom] were analyzed individually by fitting to a non-linear distortion recruitment (NLDR) model[11] to gain insights into cross-bridges mechanics (Ed, Er, b and c).

#### Myosin heavy chain composition

To determine the myosin isoform composition of muscle fibers that were used in mechanical experiments, we used Sodium dodecyl sulfate polyacrylamide gel electrophoresis as described previously.[12] In brief, after mechanical experiments, the single fibers or bundles were stored in SDS sample buffer containing 62.5 mM Tris_HCL, 2% (weight/volume) SDS, 10% (v/v) glycerol, and 0.001% (w/v) bromophenol blue at a pH of 6.8 and then were boiled for 3 minutes in 80 deg. The stacking gel contained a 4% acrylamide concentration (pH 6.7), and the separating gel contained 8% acrylamide (pH 8.7) with 30% glycerol (v/v). The gels were run for 24 h at 15 ˚C and a constant voltage of 275 V. Gels for whole muscle lysates were stained with Coomassie blue and single fiber gels were silver-stained. Gels were scanned and analyzed with ImageJ (v1.49, NIH, USA).

#### Sample preparation and gel electrophoresis

One part of the frozen biopsies was prepared as previously described.[13,14] Briefly, the tissues were grinded to fine powder with glass pestles cooled in liquid nitrogen. The powder was primed were primed at -20°C for a minimum of 20 min, then suspended in 50% urea buffer [(in mol/L) 8 urea, 2 thiourea, 0.05 Tris–HCl, 0.075 dithiothreitol with 3% SDS and 0.03% bromophenol blue pH 6.8] and 50% glycerol with protease inhibitors [(in mmol/L) 0.04 E64, 0.16 leupeptin and 0.2 PMSF] at 60°C for 10 min. Then the samples were centrifuged at 13 000 revolutions per minute (rpm) for 5 min, aliquoted and flash frozen in liquid nitrogen and stored at -80°C. Nebulin was visualized by running the solubilized samples on 1.0% vertical SDS-agarose gel [13,15] at 15 mA per gel for 3:20, staining the gel with coomassie blue, as described previously[13,16] and scanning it using a commercial scanner.

Western blots for nebulin were run with 0.8% agarose gels run for 15 mA/gel for 2 h and 50 min before being transferred to a PVDF membrane using a semi-dry transfer unit (BioRad, Hercules, CA, USA). All blots were initially stained with Ponceau S for protein visualization. Membranes were then blocked and incubated overnight at 4◦C with the appropriate primary antibodies. Both the nebulin N-terminal antibody and the SH3 antibody were provided by Dr Siegfried Labeit (Nebulin N-term 1:1000 rabbit, SH3 1:200 rabbit, University of Heidelberg, Mannheim, Germany). Secondary antibodies used were conjugated with infrared fluorophores for detection (1:20 000 goat anti-rabbit CF680, Biotium, Fremont, CA, USA and 1:20 000 goat anti-mouse CF790, Biotium). Infrared western blot was analyzed using an Odyssey CLx Imaging System (Li-Cor Biosciences, NE, USA). MHC was visualized by Ponceau S and quantified with One-D scan EX software (Scanalytics Inc., Rockville, MD, USA).

#### Thin filament length measurement

Small fiber bundles were isolated from thawed biopsies. The ends of the bundles were attached to aluminum T-clips and the solution replaced with fresh relaxing solution. Bundles were stretched ~50% of their base length. Relaxing solution was then replaced with 4% formaldehyde solution and muscles were fixed for overnight. After fixation, muscles were washed with phosphate buffer saline (PBS) and embedded in Tissue-Tek O.C.T.compound (Ted Pella Inc) and stored at -80 °C. The O.C.T. embedded specimen was sectioned into 5 µm thick (Microm HM 550; Thermo Scientific) and placed on glass slides (Fisher Scientific, size: 25X75X1 mm). Fixed tissues were permeabilized with 0.2% Triton X-100 in PBS for 20 min at room temperature on a light box to bleach out the background fluorescence, blocked with 2% bovine serum albumin (BSA) and 1% normal donkey serum in PBS for 1 hour at 4°C, and incubated overnight at 4°C with Alexa Fluor 488–conjugated phalloidin (1:1000; Invitrogen). The sections were then washed three times with PBS for 15 min and coverslips (Fisher Finest Premium Cover glass, size: 22X50X1 mm) were mounted to glass slides using Aqua Poly/Mount (Polysciences Inc.). Images were captured using a Deltavision RT system (Applied Precision) with an inverted microscope (IX70; Olympus), a ×100 objective, and a charge-coupled device camera (CoolSNAP HQ; Photometrics) using SoftWoRx 3.5.1 software (Applied Precision). The images were then deconvolved using SoftWoRx. Deconvolved images were reopened in ImageJ (http://rsb.info.nih.gov/ij), then the 1D plot profile was calculated along the myofibril direction. The plot profile was analyzed using Fityk0.9.8 (http://fityk.nieto.pl). A custom 'rectangle + 2 half Gaussian' function was used for analyzing phalloidin-stained images that consisted of a rectangle that was flanked by two half Gaussian curves. To account for actin overlapping in the Z-disk which creates a small bump in the center of the rectangle, we developed a special script designed for Fityk that de-activates the center points within the rectangle fit. This improved the subsequent fit for the 'rectangle + 2 half Gaussian' function. Thin filament length was calculated as half the width of the rectangle plus half the width of the Gaussian fit at half maximum height. SL was calculated from the distance between the centers of two adjacent Gaussian fits. We analyzed a large number of images and determined thin filament length within the SL range of 2.8–3.2 µm for all patients and controls. TFL measurement for patient 180 and one controls was performed at SL range of 3.5 -4 µm.

#### Statistical analyses

All data are represented as average ± SEM (standard error of the mean). GraphPad Prism 10.02 was used to calculate statistics. For statistical analysis one-way ANOVA, and the t-test with multiple testing corrections were used, as appropriate. To compare nebulin level, TFL or mechanical features between low and normal level nebulin patients, Student's t-test were used. To compare maximal or sub-maximal tensions between controls and each of patients, nested analysis was used. To get the relationship between nebulin level and transcript level, TFL or OM sensitivity, linear regression model was performed.
